# Cleavage of PGAM5 by the intramembrane protease PARL is governed by transmembrane helix dynamics and oligomeric state

**DOI:** 10.1101/2021.11.19.469224

**Authors:** Verena Siebert, Mara Silber, Elena Heuten, Claudia Muhle-Goll, Marius K. Lemberg

## Abstract

The intramembrane protease PARL is a crucial mitochondrial safeguard by cleaving the mitophagy regulators PINK1 and PGAM5. PGAM5 substrate determinates have not been rigorously investigated and it is unclear how uncoupling the mitochondrial membrane potential regulates its processing inversely to PINK1. Here we show that in PGAM5 several hydrophilic residues distant from the cleavage site serve as key determinant for PARL-catalyzed cleavage. NMR analysis indicates that a short N-terminal amphipathic helix, followed by a kink and a C-terminal helix harboring the scissile peptide bond, is key for a productive interaction with PARL. In difference to PINK1, PGAM5 is stably inserted into the inner mitochondrial membrane until uncoupling the membrane potential triggers its disassembly into monomers that are vulnerable to PARL-catalyzed processing. We suggest a model in which PGAM5 is a slowly processed substrate with PARL-catalyzed cleavage that is influenced by multiple hierarchical substrate features including a membrane-potential-dependent oligomeric switch.

## Introduction

The primary physiological role of mitochondria is not only producing ATP as an energy source, but also to regulate cell survival (1). Mitophagy, a selective form of autophagy, can target dysfunctional mitochondria for lysosomal degradation and protect cells from oxidative damage (2). Several regulators of mitophagy, including PINK1, Parkin and PGAM5, have been identified (3, 4). Mutations or deletions of these genes have been associated with abnormal mitophagy, which in turn has been observed in a variety of diseases, including ischemic injury, heart diseases and neurodegenerative diseases (5–7). PGAM5 belongs to highly conserved phosphoglycerate mutases and is a mitochondrial protein that lacks phosphotransferase function on phosphoglycerates, but retained activity as a serine/threonine protein phosphatase (8). Loss of PGAM5 causes accumulation of damaged mitochondria that worsen necroptosis, dopaminergic neuron degeneration, and defects in growth and cell survival, establishing a molecular link between PGAM5 and the pathogenesis of Parkinson’s disease and cardiac diseases (for review see (9)). Depending on the mitochondrial stress level, PGAM5 can either stimulate cell survival or cell death. Under mild stress, PGAM5 induces mitochondrial biogenesis and mitophagy, maintaining mitochondrial homeostasis (10, 11). Under severe stress, PGAM5 promotes mitochondrial fission and regulates multiple death signals to induce cell death (12–14). This cell death-promoting role of PGAM5 has brought the mitochondrial phosphatase into prominence for developing therapies against the above-mentioned diseases, including colon, breast and cervical cancer (15, 16). The sub-localization of PGAM5 in mitochondria is still controversial. PGAM5 contains an N-terminal non-cleaved mitochondrial targeting sequence that is also part of a transmembrane (TM) segment that anchors the C-terminal phosphatase domain to the inner mitochondrial membrane (IMM) (17, 18). Nevertheless, PGAM5 was also found to interact with several cytoplasmic proteins at the outer mitochondrial membrane, where its phosphatase domain is accessible from the cytosol (14, 19). PGAM5 is cleaved by the PINK1/PGAM5-associated rhomboid-like protease (PARL) (20), which by an ill-defined mechanism is stimulated by disruption of the inner mitochondrial membrane potential with the protonophore carbonyl cyanide m-chlorophenyl hydrazone (CCCP) (17, 21). PARL belongs to the rhomboid intramembrane proteases and was found to cleave PGAM5 in the second half of the TM domain leading to the release of the C-terminal phosphatase domain into the intermembrane space (IMS). Depending on the assay system, PARL cleavage has been mapped between amino acids F23-S24 (22) or S24-A25 (17), respectively. Recently, PARL-dependent mitochondrial release of PGAM5 that is thought to occur via proteasome-mediated rupture of the outer mitochondrial membrane through Parkin has been shown to trigger Wnt/β-catenin signaling (10,14,23). PGAM5 is known to form an equilibrium between dimeric and multimeric states (24) and catalytic activation of PGAM5 requires dodecamer formation (25, 26). Furthermore, those dodecamers can assemble into long filaments in the cytoplasm, which were described to colocalize with microtubules (25). In this process, the multimeric state of PGAM5 represents a molecular switch between mitofission/mitophagy and apoptosis. While PGAM5 multimers interact with FUNDC1 to initiate mitophagy and mitochondrial fission, PGAM5 dimers bind to Bcl-xL to prevent apoptosis (11).

Central mediator of these PGAM5 functions is PARL, which is part of the proteolytic hub formed by the iAAA-protease YME1L and the matrix scaffold protein SLP2, which is collectively known as the SPY complex (21). Interactions between the intramembrane protease PARL and the two substrates PINK1 and PGAM5 are inversely correlated: In polarized mitochondria PARL preferentially cleaves PINK1, while after mitochondrial depolarization PARL preferentially cleaves PGAM5 (17, 21). PGAM5 was described to regulate mitophagy by stabilizing PINK1 under stress conditions (18, 27). Additional and most likely simultaneous to mitochondrial protein import arrest due to disrupted membrane potential, the kinase PINK1 accumulates at the outer mitochondrial membrane where it recruits the E3 ubiquitin ligase Parkin for degradation of damaged mitochondria (28, 29). It is still unknown what the exact cleavage determinants of PGAM5 are and how the constitutive cleavage is controlled by PARL in the SPY complex. Although a consensus sequence motif around the cleavage site of a bacterial rhomboid protease substrate has been identified (30, 31), it is not entirely clear how substrate residues surrounding the cleavage site, referred to as P1 and P1’ (32), determine recognition of cognate substrate TM domains. Due to hydrophobicity of the lipid bilayer, single-spanning rhomboid substrates have to adopt a helical conformation that prevents their hydrophilic peptide backbone from contact with the membrane core (33). Substrate helices therefore have to transiently unfold near the protease active site, prior to cleavage by proteases and TM flexibility has been shown to contribute to substrate specificity of intramembrane proteases (34–37). Likewise, previous analysis of the PINK1 TM helix in cell-based assays showed that two conserved glycine residues that are predicted to lower TM helix stability are key for PARL-catalyzed cleavage (28, 38). Given the importance of PGAM5 in mitochondrial dynamics, we ask what the cleavage determinants of PGAM5 are and set out to determine these in a combination of cell-based and cell-free PARL assays with liquid-state NMR to study structural properties of the substrate TM domain.

## Results

### Phenylalanine in P1 position enables efficient PGAM5 processing by PARL but is not strictly required

Interestingly, our previous work with PINK1 showed that two glycine residues distant from the cleavage site are crucial for PARL-catalyzed cleavage (28), and recent multiplex substrate profiling indicated a preference of PARL for phenylalanine in P1 (22). However, alignment of all so far known PARL substrates does not reveal an obvious consensus sequence with many substrates including PINK1 showing other residues in P1 (Fig. S1A). Likewise, mutation of S24 in the PGAM5 cleavage site region, which when mutated to phenylalanine or tryptophan reduces PARL-catalyzed PGAM5 processing in tissue culture cells (17, 39), is not conserved across evolution (Fig. 1A). Hence, we asked whether analogous to PINK1, a less defined signature of amino acid residues enables cleavage by PARL. To this end, we expressed FLAG-tagged human PGAM5 wild type (wt) and TM domain mutants in Hek293 T-REx cells expressing a doxycycline-inducible PARL-specific shRNA (28) and analyzed processing efficiency at different PARL levels by western blotting. The uncoupler CCCP, which disrupts the inner mitochondrial membrane potential and thereby stimulates PGAM5 processing (40, 41), as well as ectopically expressed PARL were added to increase turnover of the 32-kDa full length form of PGAM5 to the processed 28-kDa species (Fig. 1B).

**Figure 1.**
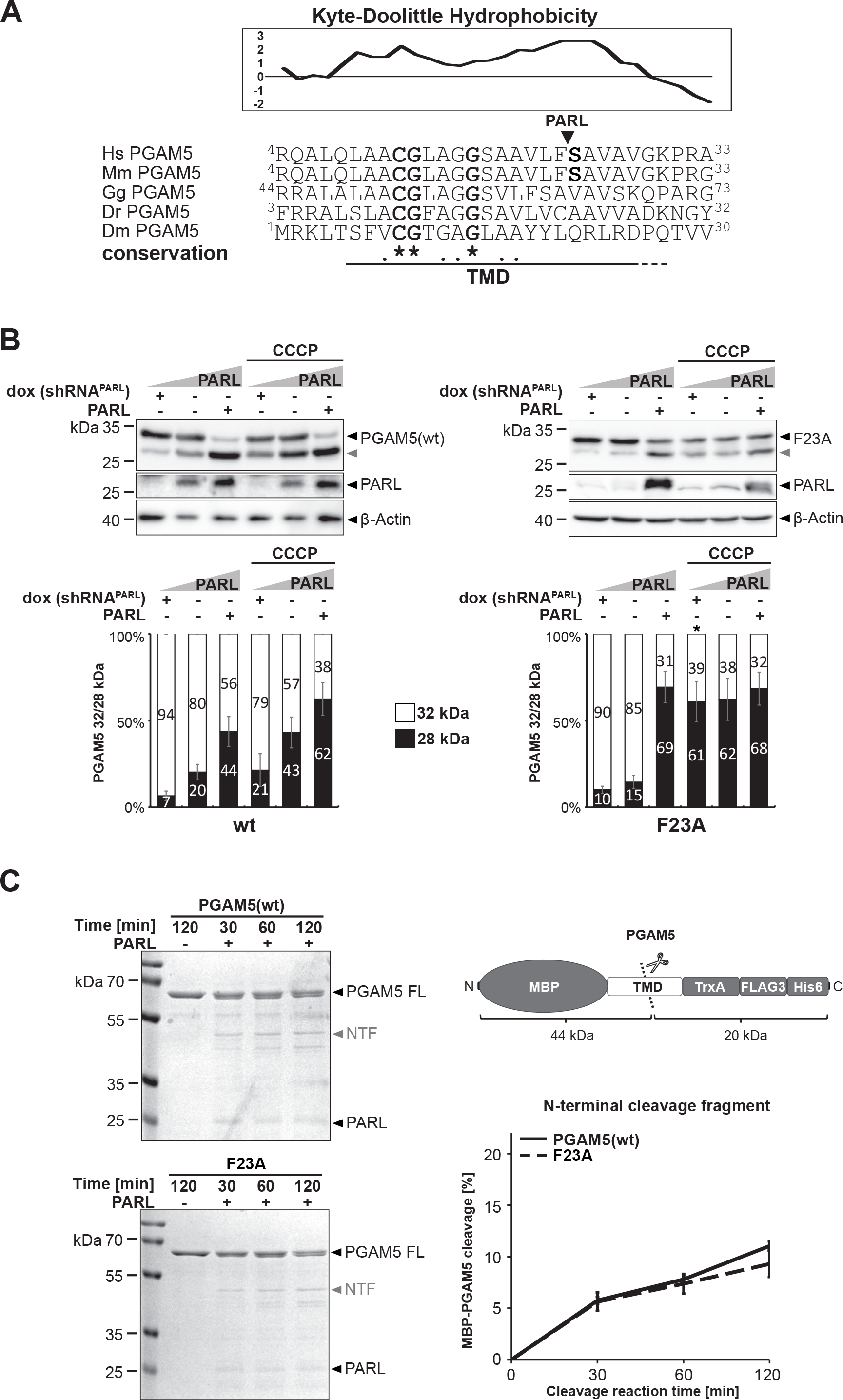
Bulky residue in P1 position shows only modest influence on PGAM5 processing. **(A)** Multi-sequence alignment reveals that C12, G13 and G17 are conserved between PGAM5 from human *Homo sapiens* (Hs), *Mus musculus* (Mm), *Gallus gallus* (Gg), *Danio rerio* (Dr) and *Drosophila melanogaster* (Dm). The predicted hydrophobic TM domain (TMD) is underlined. The hydrophobicity plot of the relevant region in human PGAM5 is shown [using the scale of Kyte and Doolittle (86) with a window size of 7] indicating the potential TMD boundaries. Arrowhead indicates the PARL cleavage site as determined by (22). **(B)** PGAM5 processing was analyzed in a cell-based PARL gain- and loss-of-function assay and western blotting. Whereas knockdown of endogenous PARL by doxycyclin (dox)-induced expression of a PARL-specific shRNA prevents PGAM5 cleavage, ectopic expression of PARL increased processing. PGAM5 processing was further stimulated by treating cells with the mitochondrial uncoupling agent CCCP. Grey arrowhead: 28 kDa cleavage fragment. Lower panel shows quantification of PGAM5 32/28 kDa distribution (n = 3, mean ± SEM). Significant changes versus wt PGAM5-FLAG are indicated with black stars (*p ≤ 0.05; unpaired two-tailed t-test). **(C)** Incubation of detergent-solubilized and purified recombinant PARL with a chimeric 64 kDa substrate containing the PGAM5 TMD (residues 1-46) fused to an N-terminal maltose binding protein (MBP) and a C-terminal Thioredoxin 1 (TrxA) domain (outlined on the right) leads to generation of an N-terminal cleavage fragment (NTF) as resolved by SDS-PAGE and staining with Coomassie blue. Of note, the 20 kDa C-terminal cleavage fragment did not become visible under the experimental conditions and therefore the N-terminal cleavage fragment was used for quantification (n = 3, means ± SEM). PARL- dependent alternative cleavage fragments appeared as side-effects of the detergent background. FL: MBP-PGAM5 full length.

Overexpression of PARL but not its catalytic-inactive mutant (PARL^S277A^) significantly enhanced levels of cleaved PGAM5 (Fig. S1B). Consistent with previous reports, knockdown of PARL prevented processing of PGAM5 wt in unstressed cells and significantly reduced generation of processed PGAM5 in presence of CCCP (17) (Fig. 1B). Additional to human tissue culture, we examined wt and mutant PGAM5 TM domains in an *in vitro* cleavage assay based on detergent-solubilized recombinant human PARL (Fig. 1C) (22). Since it is not known to what extend the amino acid sequence surrounding the scissile peptide bond influences cleavage specificity, we started analyzing the F23A mutant of PGAM5, which removes the bulky amino acid at P1 that had been shown to be favored in a peptide-based multiplex *in vitro* assay (22). In our Hek293 T-REx cell-based gain- and loss-of-function assay, we observed that at endogenous PARL level PGAM5^F23A^ is slightly less processed than PGAM5 wt but the difference does not reach significance (Fig. 1B).

Immunofluorescence microscopy analysis revealed that mitochondrial targeting of PGAM5^F23A^ was not affected by the mutation (Fig. S1C). Surprisingly, when PARL is overexpressed or the inner mitochondrial membrane potential is disrupted by CCCP, PGAM5^F23A^ gets extensively cleaved (Fig. 1B) and becomes a better substrate for the stress-activated metalloprotease OMA1 as judged by siRNA knockdown experiments (Fig. S1D).

OMA1 cleaves PINK1 and PGAM5 under certain stress-conditions and is regulated by SLP2 as part of the SPY complex (21, 42). Consistent with the cell-based PARL assay, the F23A mutant was also cleaved *in vitro* by purified detergent-solubilized PARL as efficient as the MBP-PGAM5 wt fusion protein (Fig. 1C). Taken together, these results show that a phenylalanine in the P1 position is not a strict requirement but may help to enable efficient PGAM5 processing when PARL activity is limiting. Since the PGAM5 construct with a mutated P1 position (F23A) does not show decreased cleavage but interestingly, increased cleavage under PARL overexpression and the induction of mitochondrial stress by CCCP, we suggest that additional cleavage determinants exist that dominate substrate selection.

### PARL-catalyzed cleavage of PGAM5 is influenced by multiple TM residues

In order to determine the influence of two conserved glycine and other hydrophilic amino acid residues in the TM domain of PGAM5 (Fig. 1A) on PARL-catalyzed cleavage, in a next step, we mutated them to the hydrophobic amino acid leucine or phenylalanine (Fig. 2A). Although the exact influence on TM domain stability cannot be predicted, biophysical studies in detergent micelles suggest a stabilizing effect of the helix conformation (43), which is predicted to counteract recognition of the scissile peptide bond by the rhomboid active site (34). Both single PGAM5 mutations C-terminal of the cleavage site, namely G29L and P31L, as well as a G29L/P31L double mutant did not significantly reduce PARL-catalyzed cleavage, with a tendency of G29L/P31L to a slightly reduced processing efficiency (Fig. 2B and Fig. S2A). Immunofluorescence microscopy analysis revealed that mitochondrial targeting of PGAM5 was also not affected by these mutations (Fig. S2B), indicating that the modest reduction is caused by direct effects on PARL-catalyzed processing. However, while for PINK1 mutation of a single glycine C-terminal of the cleavage site was sufficient to block processing (28), for PGAM5^G29L/P31L^ the observed reduction of PARL-catalyzed cleavage was minor only. Again, this points towards alternative cleavage determinants in the rest of the TM helix. Surprisingly, a construct with C12L mutation in the N-terminal portion of the PGAM5 TM domain is cleaved more efficiently than PGAM5 wt, whereas G13L, G16L, and G17L show decreased cleavage when compared to PGAM5 wt (Fig. 2B and Fig. S2A). Of note, immunofluorescence analysis revealed that for the G13L and G16L mutants a certain fraction is mistargeted to the Endoplasmic Reticulum (ER) (Fig. S2B). Despite showing a clear stabilization, because of the dual localization, these mutants cannot be unambiguously analyzed in cells. As it has been observed before, a S24F mutation nearly completely inhibited PARL-catalyzed processing (17) and (Fig. 2B and Fig. S2A). Taken together these results show that multiple features of the PGAM5 TM helix influence PARL-catalyzed cleavage. Strikingly, S18L was not processed, even at PARL overexpression and CCCP- stimulation (Fig. 2B and Fig. S2A) while targeting to mitochondria was not affected (Fig. S2B). However, a chimeric MBP-PGAM5 fusion with the S18L mutation in the TM domain was cleaved *in vitro* by detergent-solubilized PARL with the same efficiency as the wt construct (Fig. 2C and S2C). We speculate that the effect caused by the TM domain mutations is at least partially dependent on the context of the lipid bilayer and consequently any semi-quantitative detergent-based cleavage assay is only suitable to reveal influence of the primary amino acid sequence surrounding the cleavage site (22). Likewise, the G17L and S24F mutants, which reduced PARL-catalyzed cleavage of PGAM5 in cells and G13L, did not show striking changes in cleavage tested in DDM-micelles when compared to the wt TM domain of PGAM5 (Fig. 2C and S2C). Overall, our results indicate that PARL-catalyzed cleavage of PGAM5 is determined by multiple TM features. The strongest inhibition is observed by S18L leading to complete inhibition in the cell-based PARL gain- and loss-of-function assay. However, this residue is not conserved outside vertebrates and for example in the fruit fly *Drosophila melanogaster* a leucine residue is found at this position (Fig. 1A), which would predict that cleavage by the PARL orthologue Rhomboid-7 is hampered. Ectopic expression of FLAG-tagged *D. melanogaster* PGAM5 in human cells, which is correctly localizing to mitochondria (Fig. S2B), resulted in significantly decreased PARL-catalyzed cleavage when compared to human PGAM5 wt at endogenous PARL level (Fig. 2D).

**Figure 2.**
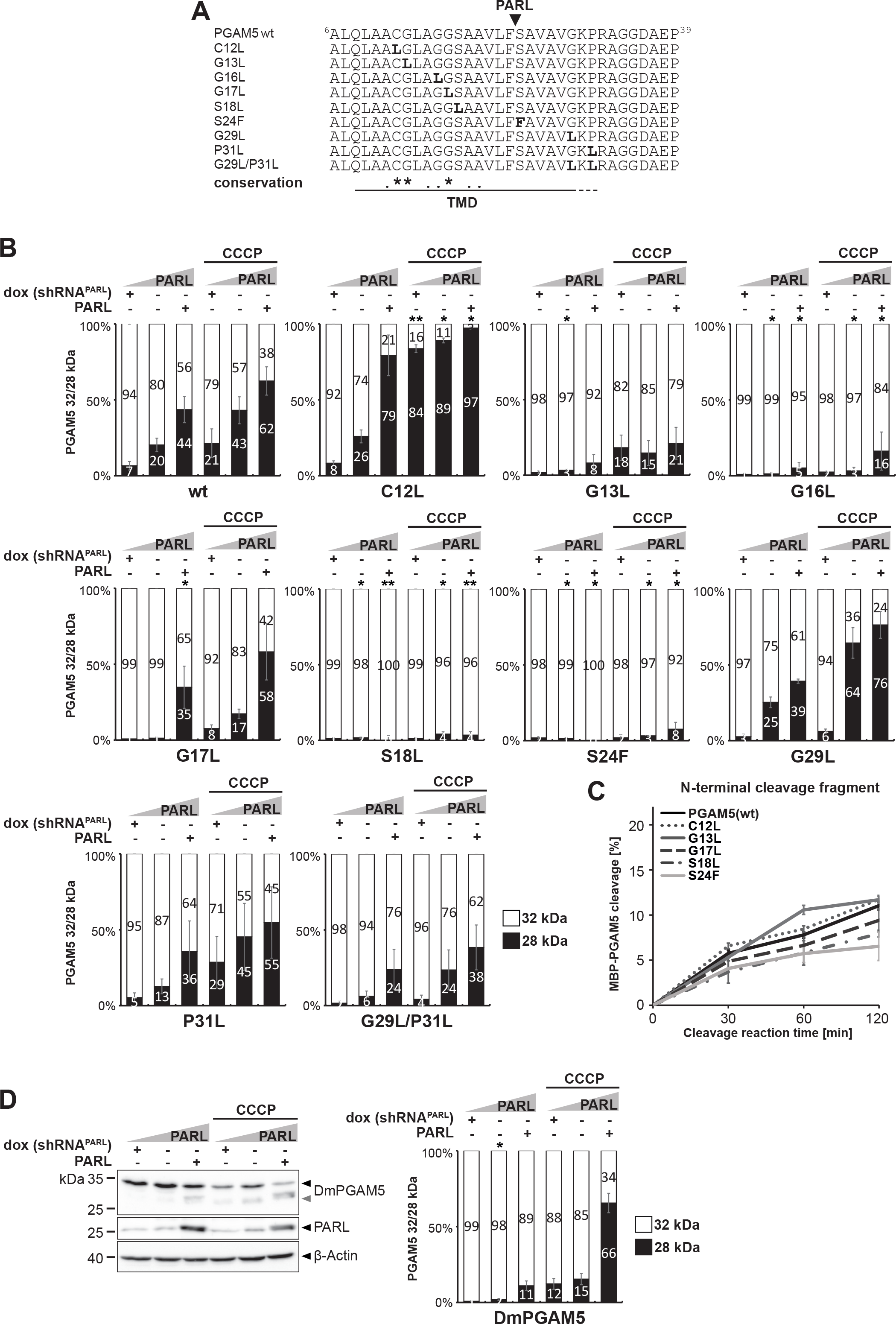
PARL-catalyzed cleavage of PGAM5 is influenced by multiple TM residues. **(A)** Amino acid sequences of TM domain mutants of human PGAM5 used in this study, including S24F previously analyzed by (17). **(B)** Quantification of PGAM5 32/28 kDa distribution upon PARL knockdown, endogenous levels or PARL over-expression without or with CCCP treatment (n = 3, means ± SEM). Significant changes versus wt PGAM5-FLAG are indicated with black stars (*p ≤ 0.05, **p ≤ 0.01; unpaired two-tailed t-test). See Fig. S2A for representative western blots. **(C)** Quantification of N-terminal cleavage fragment of purified MBP-PGAM5 as indicated (n = 3, means ± SEM; see Fig. S2C for representative Coomassie gel). **(D)** Corresponding cell-based gain- and loss-of-function assay ectopically expressing *D. melanogaster* PGAM5-FLAG with quantification of PGAM5 32/28 kDa distribution in the right panel (n = 3, means ± SEM). Grey arrowhead: 28 kDa cleavage fragment. Significant changes versus wt PGAM5-FLAG are indicated with black stars (*p ≤ 0.05; unpaired two-tailed t-test).

However, processing efficiency was higher when compared to the S18L mutant of human PGAM5 (Fig. 2B), indicating that the inhibiting property of leucine can be balanced by compensatory changes such as additional charged TM residues in *D. melanogaster* PGAM5, namely R22 and R24 (Fig. 1A). However, the length of the TM region is reduced by 4-5 residues in *D. melanogaster* as well as in *Aedes aegypti* (yellow fever mosquito) and *Caenorhabditis elegans* (nematode), leaving it elusive which amino acid residues at certain positions are essential for cleavage by rhomboid proteases across the animal kingdom.

While for most PGAM5 TM residues there seems to be no strong selective pressure in evolution, C12 is shared between various species in addition to G13 and G17 (Fig. 1A), albeit not to 100%. Among vertebrates, for instance, *Xenopus laevis* (African clawed frog) and *Bufo bufo* (common toad) do not contain a cysteine at this position and neither do *A. aegypti* or *C. elegans*. As mutation of C12 to leucine caused an unexpected increase of PARL-catalyzed cleavage of human PGAM5 in our cell-based assay (Fig. 2B and Fig. S2A), we further investigated its role in substrate selection by mutating it to a serine (Fig. 3A), which is more hydrophilic than leucine and closer to the chemical properties of cysteine.

**Figure 3.**
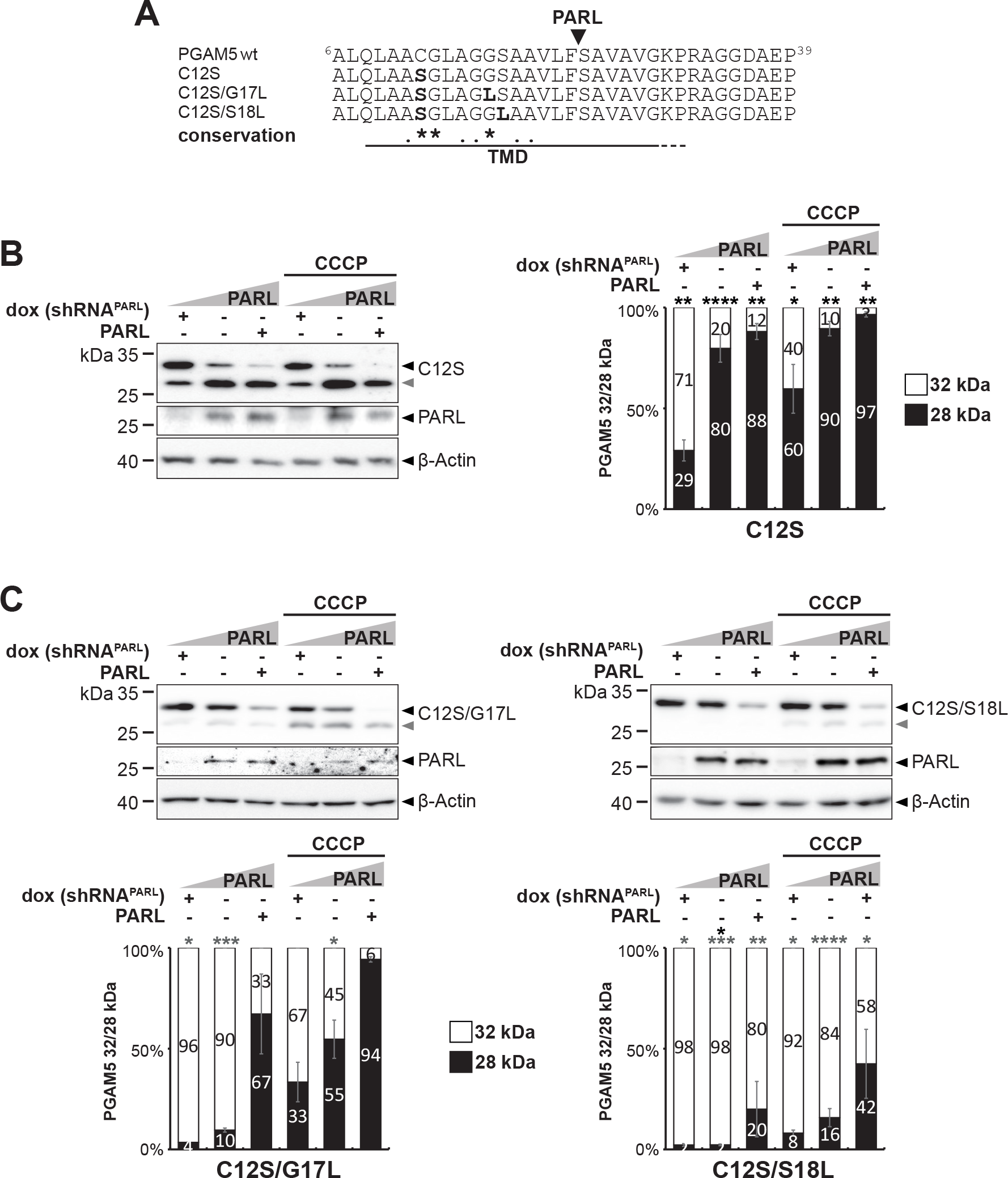
N-terminal substrate feature in PGAM5 important for PARL-catalyzed cleavage. **(A)** Amino acid sequences of PGAM5 C12S TM domain mutants used in this study. **(B)** Mutation of C12 to serine further increases processing efficiency in cell-based PARL gain- and loss-of function assay as in Fig. 1B. Grey arrowhead: 28 kDa cleavage fragment. Right panel shows the quantification of PGAM5 32/28 kDa distribution (n = 3, means ± SEM). Significant changes versus wt PGAM5-FLAG are indicated with black stars (*p ≤ 0.05, **p ≤ 0.01, ****p ≤ 0.0001; unpaired two-tailed t-test). **(C)** C12 acts independent of G17 and S18. Double mutants with the C12S mutation show decreased cleavage efficiency. Grey arrowhead: 28 kDa cleavage fragment. Lower panel shows quantification of PGAM5 32/28 kDa distribution upon PARL knockdown, endogenous levels or PARL over-expression without or with CCCP treatment (n = 3, means ± SEM). Significant changes versus wt PGAM5-FLAG are indicated with black stars, significant changes versus PGAM5^C12S^-FLAG are indicated with grey stars (*p ≤ 0.05, **p ≤ 0.01, ***p ≤ 0.001, ****p ≤ 0.0001; unpaired two-tailed t-test).

C12S was correctly targeted to mitochondria (Fig. S3A) and interestingly, this mutation even further increased processing significantly, especially at endogenous PARL level (Fig. 3B) when compared to C12L (Fig. 2B and Fig. S2A). The enhanced cleavage was confirmed to be not induced by OMA1 activity based on siRNA knockdown experiments (Fig. S3B-C).

However, when combined with the processing-inhibiting G17L or S18L mutations, the double mutants C12S/G17L and C12S/S18L showed significantly decreased cleavage efficiency when compared to C12S (Fig. 3C), indicating that the substrate features act independently and show additive effects. Mutation of C12, which is 11 amino acids away from the PARL cleavage site, might help to render the TM domain into the PARL active site and thereby increase cleavage efficiency. Interestingly, only the C12S TM domain mutant showed a slightly increased cleavage in the *in vitro* PARL assay when compared to wt MBP-PGAM5 (Fig. S3D). Taken together with the cell-based assay, our mutagenesis of hydrophilic TM residues revealed that PARL-catalyzed cleavage of human PGAM5 is influenced both by TM residues N-terminal and C-terminal of the scissile peptide bond. For this long-range influence, especially cysteine-12 plays a prominent role.

### Structural properties of the PGAM5 TM domain

To understand whether the different cleavage efficiencies observed for PGAM5 mutants are caused by structural or dynamic effects, we determined the structure of PGAM5 wt and mutant TM domains (residue 2 to 35). To this end, we used aqueous trifluoroethanol (TFE) as a model that is believed to mimic the biophysical properties of a water-filled intramembrane protease active site cavity (44, 45). Circular dichroism (CD)-spectroscopy revealed that all TM domains showed a moderate content of α-helical structure in the range of 33-38% in this solvent (Fig. S4A), indicating that it is a suitable model situation to study unfolding of the PARL substrate TM helix. Mutation of the central glycine G17 to a hydrophobic leucine slightly increased helicity with respect to wt, whereas mutation of C12 to either serine or leucine did not result in explicit secondary structure changes. NMR secondary chemical shifts are sensitive reporters of secondary structure. They are calculated as difference between measured Hα or Cα chemical shifts and the respective chemical shifts in random coil peptides (46). Fig. 4A shows that in the model situation of TFE/water the PGAM5 TM domain is divided into two distinct α-helical parts R4-C12, N-terminally three residues longer than the predicted TM part, and S18-V28 with negative Hα and positive Cα secondary chemical shifts. The central part, G13-G17, had no preference for a defined secondary structure, generating a hinge-like loop. This NMR analysis in a model situation suggests that the PGAM5 TM domain is not a straight, single helix but instead shows a kink in the region of the PARL active site splitting it into two helices with the longer, C-terminal end harboring the scissile peptide bond.

**Figure 4.**
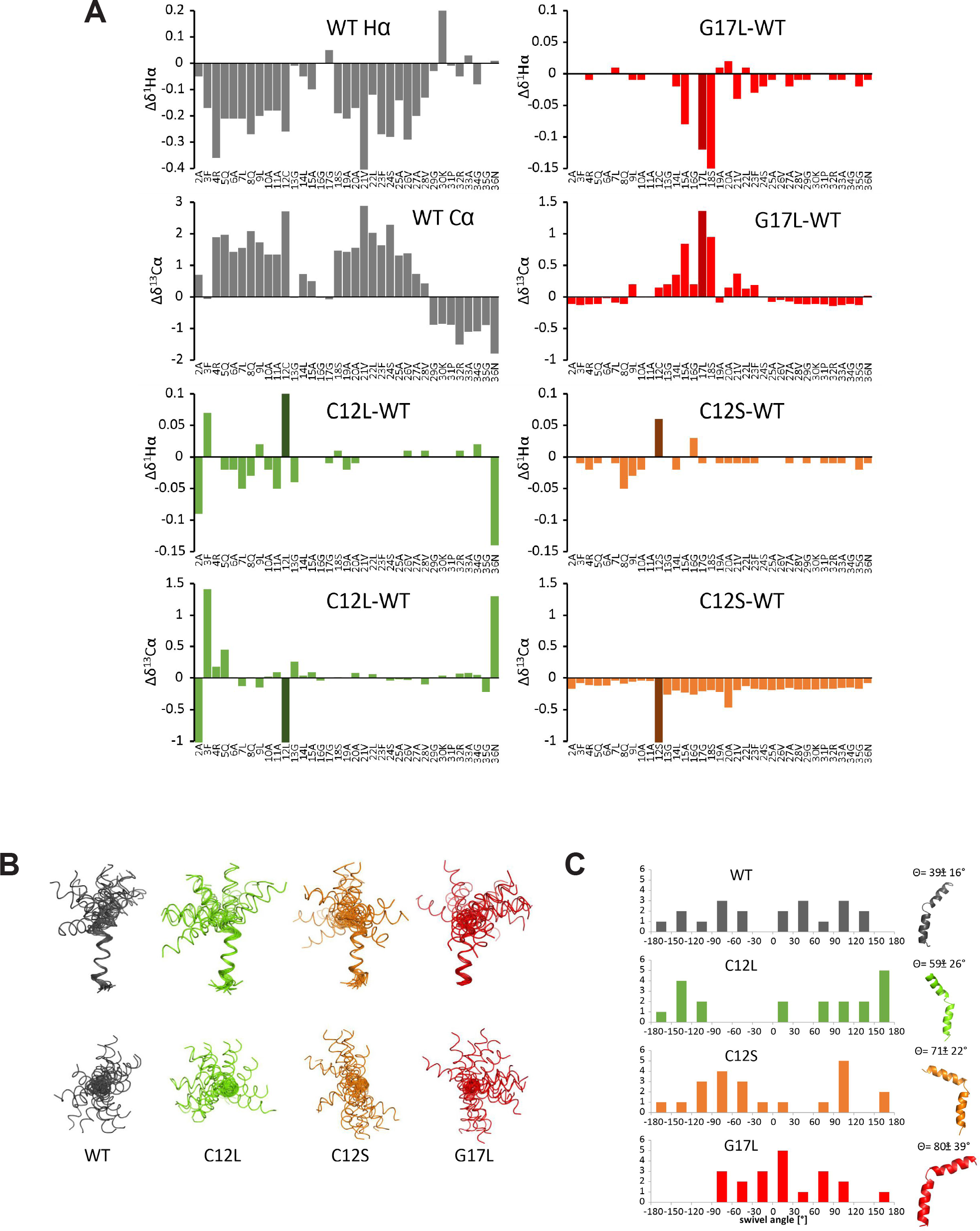
Strucutral properties of the PGAM5 TM domain. **(A)** Random coil chemical shifts were subtracted from experimental values of Hα and Cα respectively. Negative secondary chemical shifts of Hα and positive secondary shifts of Cα indicate α-helical structure. For C12L, C12S and G17L deviations from wild type secondary chemical shifts are shown. Negative values for Hα and positive values for Cα suggest a more helical structure compared to wild type. **(B)** Upper panel front view, lower panel top view. All structures aligned from residue 20 to 25, L22Hα defined as x-axis. Black wt, green C12L, orange C12S, red: G17L. **(C)** The swivel angle is defined by the rotation of the N-terminal helix relative to the Hα atom of L22 as reference in the C-terminal helix. Swivel angles of the 20 best structures were grouped in 30° segments, frequency distributions are given above. Right: The bend angle is defined as the angle between the axis through the N-terminal and the C-terminal helix. Bend angles and representative structures are given above.

In order to analyze stability of this unusual TM domain, we studied which amide protons were protected against deuterium exchange by recording short consecutive ^1^H^1^H-TOCSY experiments and following the intensities of the HN-Hα crosspeaks. H/D exchange monitored this way probes for stable hydrogen bonds (Fig. S4B). Although the exchange of several residues could not be determined due to spectral overlap, two regions in PGAM5 TM domain could be marked that showed reduced deuterium exchange. Slowed down exchange in Q8- C12 in the N-terminal helix and A19-V28 in the C-terminal helix was interrupted by the region G13-G17 showing immediate exchange without involvement in stable hydrogen bonds. This corroborates our analysis of secondary chemical shifts that this region has no defined secondary structure and may serve as a hinge. However, mutants C12S and C12L only marginally affected secondary structure, because chemical shift changes were small and dispersed over the entire TM domain. Mutation of C12 to leucine showed disturbances within the N-terminal helix that cannot be easily interpreted in terms of secondary structure changes. Mutation to serine seemed to slightly destabilize the entire TM helix. G17L seemed to induce α-helical structure in the central part G13-L17 with strong alterations in both Hα and Cα secondary chemical shifts. Since changes in secondary structure caused by the mutants were in total inconspicuous, we calculated their 3D structures (Table S1). Fig. 4B shows the bundle of the 20 best structures each, superposed onto the C-terminal helix. The extent of either N- or C-terminal helix did not vary between the four TM peptides and no further major structural changes could be discerned. This was intriguing with respect to the observed changes in cleavage efficiency and taking the TFE/water model into account we ruled out simple local structure changes as facile explanations. The superposition showed that the orientation of the N-terminal helix with respect to the C-terminal one was not well defined for all four bundles. We wondered whether the orientation was fully arbitrary or whether certain conformations were preferred. Looking at the bundle from the top when the C-terminal helix was aligned along the –z axis, the wt fanned out into two possible conformation ranges where two angle ranges of ∼ 60° each were devoid of structures (Fig. 4C). Interestingly, the two mutants, which are more readily cleaved, C12S and C12L, showed also restricted conformational variability. The angle region devoid of structures was here, however, much more pronounced apart for two structures in C12L. The area of possible conformations of C12S overlapped with one of the conformational regions in the wt whereas the bundle in C12L was turned by roughly 90°. G17L on the contrary stabilizes the beginning of the C-terminal helix elongating it on one hand and restricting the possible mutual orientations of the two helical parts. G17L, the mutant where cleavage efficiency dropped considerably, had a distribution of possible orientations that was distinct from the other three by roughly 120°.

This was caused by the slight elongation of the C-terminal helix. Taken together these results in the TFE/water model indicate that the N-terminal feature in PGAM5’s TM domain affects TM substrate dynamics and thereby may enable or hamper bending into the PARL active site. However, in the absence of structural data in the lipid bilayer of the PGAM5 TM domain and PARL we note that this remains speculative.

### Formation of the PGAM5 higher order structure prevents PARL-catalyzed cleavage

In addition to its TM domain, PARL-catalyzed cleavage of PGAM5 may be influenced by its C-terminal portion facing the IMS. While a negatively charged motif C-terminally to the TM anchor of PINK1 and STARD7 facilitates PARL-catalyzed cleavage (42, 47), for PGAM5 such a pronounced cluster of negatively charged amino acids cannot be found at the same position (Fig. S5A). We therefore asked whether introducing negative charges to the PGAM5 ‘juxtamembrane region’ might increase PARL-catalyzed processing as well. Replacing two glycine residues C-terminal of the TM helix by glutamic acid did not change processing in unstressed cells, but under the CCCP treatment conditions PGAM5^GG34/35EE^ was significantly more cleaved when compared to PGAM5 wt (Fig. 5), while correctly localizing to mitochondria (Fig. S5B). Control experiments under *OMA1* knockdown confirmed that the processing is catalyzed by PARL and no significant role of OMA1 activity was observed (Fig. S5C). From these observations we conclude that substrates like PINK1 or STARD7 can be seen as ‘fast’ processing substrates, whereas PGAM5 is lacking the advantageous negative charges and may be processed in unstressed mitochondria by PARL with a slower kinetic.

**Figure 5.**
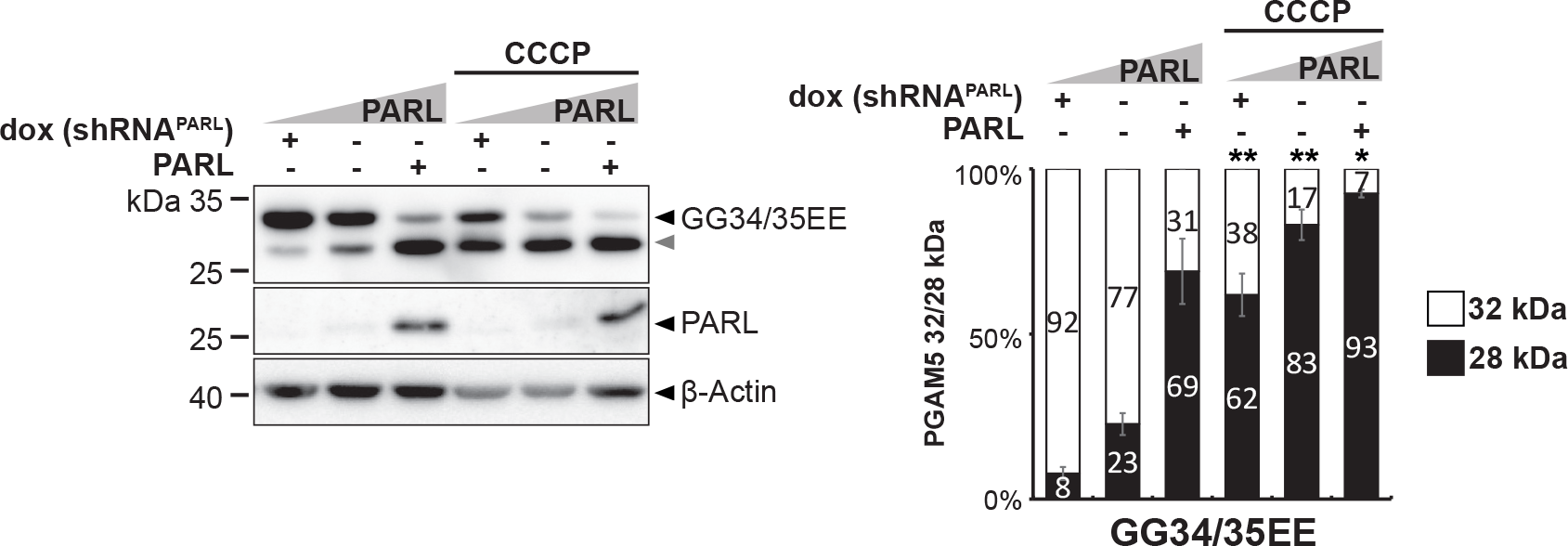
Negative charges in the PGAM5 juxtamembrane region influence cleavage efficiency under CCCP. Processing of PGAM5 mutant with negative charged juxtamembrane region was analyzed in a cell-based PARL gain- and loss-of-function assay as in Fig. 1B. Grey arrowhead: 28 kDa cleavage fragment. Right panel shows quantification of PGAM5 32/28 kDa distribution (n = 3, means ± SEM). Significant changes versus wt PGAM5-FLAG are indicated with black stars (*p ≤ 0.05, **p ≤ 0.01; unpaired two-tailed t-test).

As PGAM5 is known to form oligomers, slower processing speed may allow PGAM5 imported into mitochondria to form higher molecular assemblies. Because intramembrane proteases, such as γ-secretase, are commonly thought to cleave their substrates only in a monomeric state (48–51), we asked whether PGAM5 processing is affected by its higher order structure. Hence, we tested a monomeric PGAM5 mutant lacking its C-terminal dimerization domain (ΔC) and a multimerization-deficient mutant lacking the WDxxWD-motif (AAxxAA) (Fig. 6A) (24) in our cell-based PARL gain- and loss-of-function assay.

**Figure 6.**
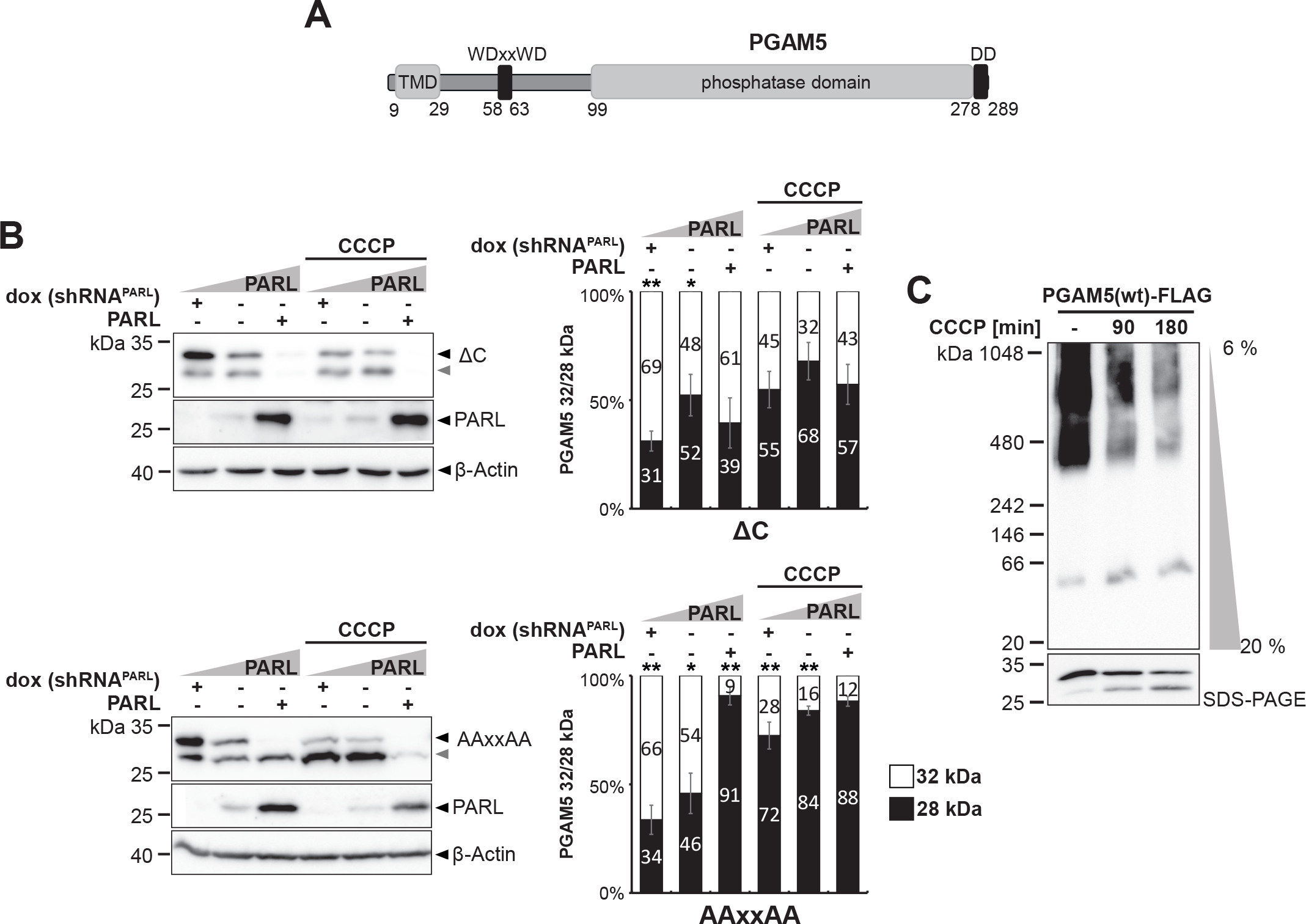
Formation of the PGAM5 higher order structure prevents PARL-catalyzed cleavage. **(A)** Schematic representation of PGAM5 domain structure indicating TM domain (TMD), WDxxWD multimerization motif, and C-terminel dimerization domain (DD). **(B)** Processing of monomeric PGAM5^ΔC^ (Δ278-289) and multimerization-deficient PGAM5^AAxxAA^ was analyzed in a cell-based PARL gain- and loss-of-function assay as in Fig. 1B. Grey arrowhead: 28 kDa cleavage fragment. Right panel shows quantification of PGAM5 32/28 kDa distribution (n = 3, means ± SEM). Significant changes versus wt PGAM5-FLAG are indicated with black stars (*p ≤ 0.05, **p ≤ 0.01; unpaired two-tailed t-test). **(C)** Analysis of PGAM5 higher molecular weight structures in BN-PAGE upon treatment with 10 µM CCCP for 90 min and 180 min.

Immunofluorescence microscopy analysis revealed that mitochondrial targeting of these PGAM5 constructs was not affected by the mutations (Fig. S6A). Strikingly, PGAM5^ΔC^ as well as PGAM5^AAxxAA^ were significantly more processed by PARL when compared to PGAM5 wt, which occurred even in the absence of CCCP (Fig. 6B). Thus, cleavage of these mutants seems to be uncoupled from the physiological activation mechanism. Further increase of cleavage could be induced by additional ectopic expression of PARL and treatment with CCCP. Control experiments under *OMA1* knockdown confirmed PARL-catalyzed cleavage and no significant role of OMA1 activity (Fig. S6B). In contrast, combining the deletion of the dimerization domain (ΔC) with the G17L and S18L TM mutations significantly decreased cleavage efficiency for the double mutants when compared to PGAM5^ΔC^ alone (Fig. S6C).

This observation suggests that oligomeric state influences PARL-catalyzed processing independent of the determinants within the TM domain. Next, we asked whether CCCP may increase PGAM5 processing by disassembling its oligomers, thereby making PGAM5 monomers susceptible for PARL-catalyzed cleavage. Analysis of PGAM5 ectopically expressed in Hek293T cells untreated and treated with CCCP by blue-native (BN)-PAGE revealed a reduction of higher molecular weight assemblies in the range of 500 kDa over time of CCCP treatment (Fig. 6C). Consistent with a link to PARL-catalyzed cleavage, we observed an increase of monomeric and processed PGAM5 by BN-PAGE and SDS-PAGE. Taken together, these results reveal that PGAM5 processing is governed by an oligomeric switch that in healthy mitochondria prevents PARL-catalyzed cleavage and enables the conversion of higher molecular weight assemblies to its soluble form upon stress-induced disassembly, resulting in subsequent cleavage because of a suitable TM domain.

## Discussion

In this study, we investigated the requirements for PARL-catalyzed PGAM5 cleavage to further understand how its cleavage is accelerated by uncoupling the mitochondrial oxidative phosphorylation and thereby disrupting the mitochondrial membrane potential with the protonophore CCCP. We showed that the N-terminal portion of PGAM5’s TM domain is a critical determinant for processing by PARL. Interestingly, besides cleavage resistant forms, we obtained PGAM5 mutants that were better cleaved by PARL uncoupling it from its native regulation. Moreover, we found that a balanced net charge of the PGAM5 C-terminal juxtamembrane region prevents its premature processing by PARL allowing assembly of cleavage-resistant PGAM5 oligomers upon mitochondrial import. In contrast, we propose a model in which CCCP uncoupling the membrane potential at the IMM disassembles PGAM5 by an unknown mechanism into monomers that are efficiently cleaved by PARL to trigger PGAM5’s downstream activities. Taken together, our findings indicate that the substrate recognition mechanism of PARL depends on multiple hierarchical substrate features including a membrane-potential-dependent oligomeric switch.

### Is intramembrane cleavage of PGAM5 affected by TM helix dynamics?

Proteolytic cleavage within a TM domain is mechanistically more complex than proteolysis within an aqueous environment (34). In addition to limited availability of water, restricted lateral diffusion of the substrate and its inability to freely rotate within the lipid bilayer introduce several additional constraints. Consequently, enzyme-substrate interaction of rhomboid proteases and subsequent intramembrane cleavage is seen as a multi-step process. Prior to cleavage, the scissile peptide of the substrate has to bind into a water-filled catalytic cleft, which requires translocation of the helical substrate TM domain from the lipid bilayer towards the rhomboid protease active site. It is commonly believed that the TM helix of rhomboid substrates initially dock onto a membrane-integral exosite of the enzyme, a process that may be associated with structure-encoded global motions of the substrate TM helix (52). Subsequent unwinding of the bound TM helix allows access of the catalytic residues to the cognate cleavage site motif, followed by processing of the substrate (31, 53). In the case of bacterial and eukaryotic secretory pathway rhomboids, like GlpG and human plasma membrane rhomboid RHBDL2, substrate cleavage sites map at the N-terminal TM domain boundary and processing efficiency is largely determined by the primary sequence (52,54,55). Hence, cleavage sites are likely to access the catalytic center from the top of the enzyme (facing the outside of the cell), which demands substrate unfolding between the scissile peptide bond and the hydrophobic TM helix and a sharp turn in the protein main chain (56) while the TM helix may remain bound to the exosite (52, 56).

Since PARL is predicted to have an inverted active site (facing the mitochondrial matrix) compared to bacterial and secretory pathway rhomboids (with an outwards orientation) (57) and cleaves its canonical substrates towards the C-terminal portion of their TM domains (Fig. S1A), TM helix unwinding may play a more prominent role. Consistent with this, we now show that the preference of bulky amino acids in the P1 position (22) only results in modest effects and cleavage rate may be primary governed by TM helix dynamics. For PINK1 conserved helix-destabilizing glycine residues in the C-terminal portion of the TM domain are invariant for PARL-catalyzed cleavage (28). Substitution of equivalent putative helix-destabilizing residues in PGAM5 (G29 and P31) did only moderately impact on PARL-catalyzed cleavage, probably because these residues are located outside the helical region according to our structures, and the critical residues were found in the N-terminal half of the substrate TM domain. This suggests that TM domain dynamics are influenced by multiple features, making it difficult to predict. Using a TFE/water model system for NMR analysis that mimics important biophysical aspects of an intramembrane protease active site (44, 45), we observed no significant secondary structure changes for the mutants C12S, C12L, and G17L. Given the striking differences in the efficiency of PARL-catalyzed cleavage of these mutants observed in cells, this finding was surprising and suggests that not primarily TM helix stability determines cleavage rate. Studying the structure in the TFE/water system, we revealed that PGAM5 has a pronounced loop of five residues at the center of its TM domain between G13 and G17, several residues apart from the scissile peptide bond leading to a kink in the presumed TM part. A deviation from a straight TM helix is also observed in other intramembrane protease substrates, for example mammalian APP-C99 with a double-glycine hinge (58). Bacterial TatA (59), which is cleaved by the rhomboid protease AarA in *Providencia stuartii,* shows an even more pronounced kink in the protein main chain compared to APP-C99. This leads to an almost rectangular arrangement of the TM domain and the following amphipathic helix (60). Glycine and proline were shown to have the strongest destabilizing effects of all amino acids on model TM helices with regard to their helicity in detergent micelles (61, 62) and glycine was found twice as abundant in TM helices than in water-soluble helices (63), highlighting its importance in the functional role of TM domains. The hinge region of the *P. stuartii* rhomboid substrate TatA is formed by glycine-serine-proline, whereas the APP-C99 hinge displays a di-glycine sequence (64). Also, for PGAM5 glycine seems to comprise a major role as the hinge is formed between G13 and G17, containing a di-glycine motif with G16 and G17 (Fig. 1A). Recently, it could be shown that modulation of the hinge flexibility in the TM domain of APP-C99 alters γ-secretase cleavage (65–67) and affects substrate-enzyme interaction (68). Since the G13L, G16L, and G17L mutants of PGAM5 showed decreased cleavage when compared to PGAM5 wt in tissue culture cells (Fig. 2B, S2A), we speculate that the substrate-enzyme interaction became negatively affected within the membrane, as seen for APP-C99 and γ-secretase. The N-terminal helix of PGAM5’s TM domain from R4 to C12 shows signs of amphipathicity with R4, Q5, Q8, and C12 aligned on one side of the helix while the C-terminal helix from S18 to V29 has a strong hydrophobic character. In the TFE/water model, both helices are bent by more than 30° regarding each other leading to a putative submerged orientation of the amphipathic N-terminal helix in lipid bilayers and a strong tilt with respect to the membrane normal. The inverted topology of PARL does not allow easy extrapolation from structural details observed in the *E. coli* GlpG crystal structure. We used the model of PARL generated by AlphaFold (69) entry Q96HS1 (Fig. S7) to study similarities and differences to GlpG. Like GlpG the catalytic S277 and H335 face a water-filled cavity that in the case of PARL opens to the matrix. However, whereas GlpG cuts within the N-terminal unfolded region adjacent to the TM helix, PARL cuts within the most stable part of the TM domain close to the C-terminal end of the TM region. Our model (Fig. S7) may indicate how the PGAM5 TM domain binds to a putative PARL exosite. Insertion depth into the inner mitochondrial membrane was determined with the OPM server (70) both for the PARL AlphaFold model as well as for the PGAM5 TM domain. While in this speculative model the cleavage site (F23-S24) located in the C-terminal helix of the PGAM5 TM domain would be positioned outside the water-filled cavity, it is attractive to speculate that upon unfolding of the N-terminal helix into the matrix it may fit into the catalytic cleft. Interestingly, different swivel angles and thus different orientations of the PGAM5 N-terminal amphipathic helices could be observed in our NMR analysis. If mutant PGAM5 TM domains show these differential bending capacities also in biological membranes upon binding to the putative PARL exosite, altered directions of the cone opening might influence the cleavage efficiency. One possible scenario is that the long hinge-like loop formed by G13-G17 may allow the N-terminal amphipathic helix to swing into contact with the enzyme and that this motion is disturbed by the mutations. We are aware though that the TFE/water system is a technical compromise and in future it will be interesting to study the same mutants reconstituted in bicelles, multilamellar vesicles or proteoliposomes to get further insights into the structural and dynamic properties of PGAM5.

### Negatively charged juxtamembrane region accelerates PARL-catalyzed cleavage

In addition to TM domain properties, recognition of intramembrane protease substrates is also influenced by substrate features outside the membrane. Likewise, the yeast PARL orthologue Pcp1/Rbd1 recognizes a stretch of negatively charged amino acids located C-terminally to the cleavage site in the IMS region of its substrate Mgm1 (71). A similar negatively charged patch was suggested to influence the fate of PARL substrates such as PINK1 and STARD7 (47), while it is missing in PGAM5. In the case of PINK1 the negatively charged cluster is required for PINK1 import arrest, recognition and subsequent cleavage of the mitochondrial import intermediate by PARL. As recently published, mutant PINK1^3EA^ lacking this motif fails to accumulate on depolarized mitochondria, gets constantly imported, is interfering with the biological equilibrium and thus becomes a substrate of the stress-activated metalloprotease OMA1 (42). Here, we show that introducing negative charges into the juxtamembrane region of PGAM5 correlates with enhanced CCCP-induced PARL cleavage. Thus, we speculate that a negatively-charged C-terminal juxtamembrane region can serve as an additional cleavage determinant of PGAM5, as it may facilitate binding to a putative IMS-exposed PARL exosite.

### PGAM5 multimerization prevents processing

The intramembrane protease γ-secretase is a multi-subunit protease complex (72) and has been shown to cleave its substrates only in a monomeric state (50, 51). It is believed that TM domain dimerization, like in the γ-secretase substrate APP-C99, restricts transition into the active site, which is gated by the γ-secretase complex partner Nicastrin (73–75). PARL is also embedded in a multiprotein assembly known as SPY complex (21), and substrate gating may be similarly controlled and influenced by the oligomeric state of its substrates. PGAM5 can be found in an equilibrium between dimeric and multimeric states (24, 25), depending on its biological function as result of mitochondrial quality control. So far, the impact of PGAM5 oligomeric state on PARL catalyzed cleavage has not been addressed yet. In this work, we reveal that PGAM5 processing is affected by its oligomeric state, which potentially acts as an oligomeric switch that in response to mitochondrial stress enables recognition and conditional cleavage of PGAM5 by PARL as has been observed before (17). Thereby, PARL-catalyzed processing of the monomeric form of PGAM5 shows parallels to other rhomboid family proteins in protein quality control, which is exemplified by the ER-associated degradation pathway that removes orphan subunits of multiprotein complexes (76). It will be interesting to reveal whether PARL has a more general role in the control of inner membrane protein complexes and to decipher the molecular mechanism of how the inner mitochondrial membrane potential or general mitochondrial stress affects the oligomeric state of PGAM5.

### Model of PARL-catalyzed PGAM5 cleavage in comparison to PINK1

Depending on the stress level and in an inversely correlated manner, PARL cleaves PINK1 in healthy mitochondria as an import intermediate and PGAM5 in damaged mitochondria with a disrupted inner membrane potential as fully imported protein (Fig. 7). We hypothesize that primarily the speed of processing determines this different outcome. Because of a negative charged cluster in its juxtamembrane region and suitable TM helix, PINK1 is rapidly processed as import intermediate leading to constant release of the C-terminal cleavage fragment into the cytoplasm (77). In contrast, PGAM5 can be seen as slowly processed substrate that is inserted into the IMM as homodimer or even in a multimeric state, which withstands cleavage by PARL. This allows PGAM5 to persist in its membrane anchored form until IMM depolarization or other forms of mitochondrial stress trigger its disassembly into monomers that become subject for PARL-catalyzed cleavage (Fig. 7). In contrast, the PGAM5 mutants C12S and C12L are more efficiently cleaved by PARL in absence of the uncoupler CCCP, suggesting that they might be cleaved before they can dimerize. Hence, like for other cellular proteins monomeric forms of IMM proteins are more vulnerable to cleavage and degradation (78), which in some cases may be used in terms of quality control in order to remove orphan subunits of multiprotein complexes. For PGAM5, this dynamic detachment from its membrane anchor and a subsequent release into the cytoplasm by an ill-defined mechanism increases the range of actions from control of mitophagy to Wnt signaling (10,13,14,17,21,79). Cytosolic PGAM5 can further assemble into symmetric rings, which can further polymerize into filaments that were described to colocalize with microtubules (24, 25). Whether this phenomenon links PGAM5 to stress-induced retrograde trafficking of mitochondria or if PGAM5 filaments are initially generated inside the mitochondria is unknown and needs further investigation. In our study, we observed PGAM5 mutants that were processed stronger than PGAM5 wt but still behaved different than PINK1. This reveals that the fate of PGAM5 and PINK1 is determined by multiple factors. Given the importance of PGAM5 in mitochondrial dynamics, our foundational research on requirements for PARL-catalyzed cleavage of PGAM5 contributes to a multifaceted understanding of disease-promoting mechanisms.

**Figure 7.**
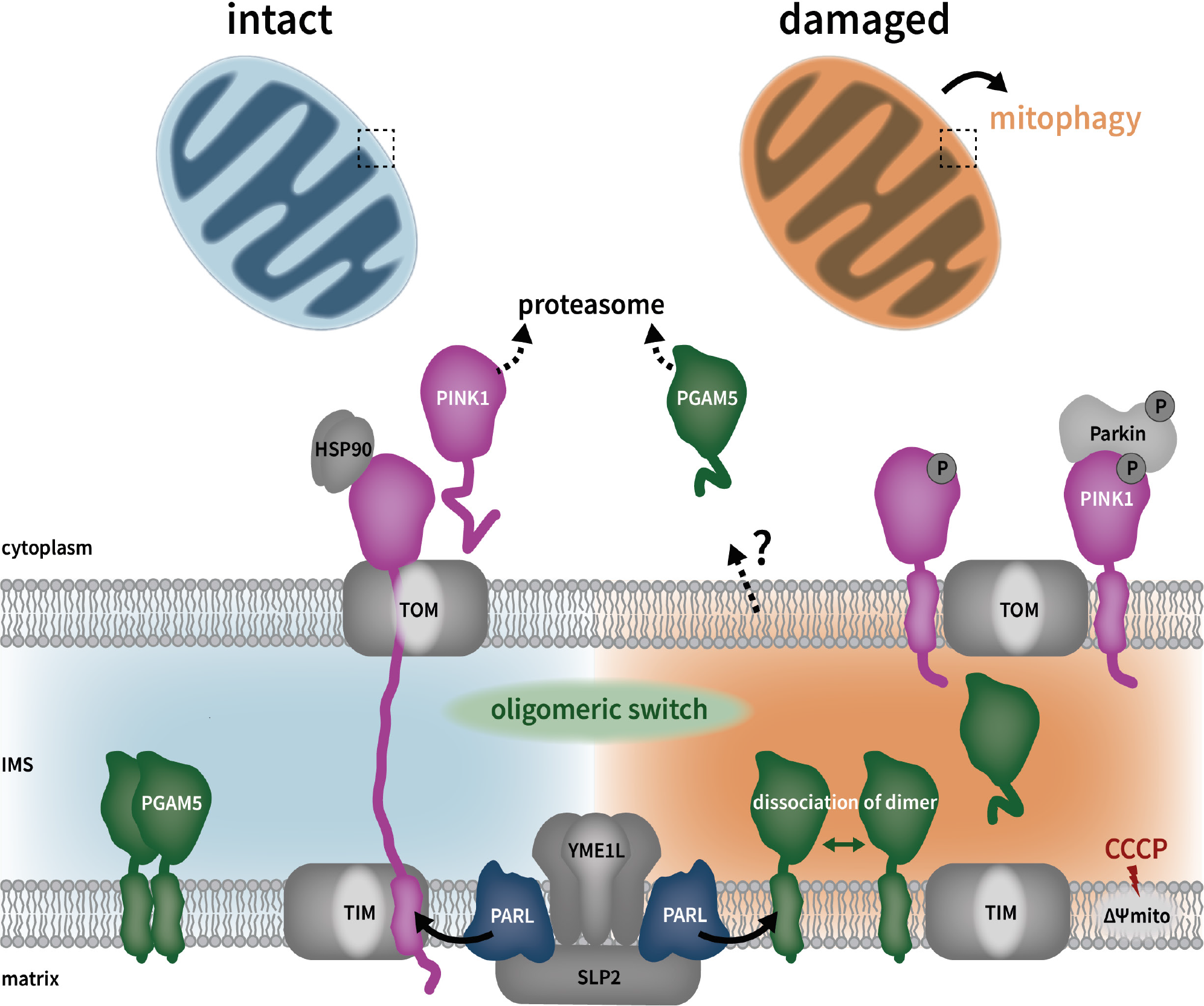
Model of PGAM5 cleavage in comparison to PINK1. Depending on mitochondrial stress, PARL cleaves PINK1 (as an import intermediate) or PGAM5 (as fully imported protein) in an inversely correlated manner. Upon disruption of the mitochondrial inner membrane potential (ΔΨmito), PGAM5 dimers or even oligomers disassemble into monomeric forms representing an “oligomeric switch” before getting processed by PARL. A portion of cleaved PGAM5 is released into the IMS, while another portion is released via a so far unknown mechanism into the cytoplasm where it undergoes proteasomal degradation.

## Materials & Methods

### Plasmids

Construction of pcDNA3.1-PARL, pcDNA3.1-PGAM5-FLAG, pcDNA3.1-PGAM5^S24F^-FLAG and recombinant pET25b(+)-MBP-PGAM5 expression plasmids have been described previously (17,22,28). Mutations in the TM domain, WDxxWD motif and juxtamembrane region of PGAM5 were introduced by Quik-Change site-directed mutagenesis (Stratagene). *D. melanogaster* PGAM5 was ordered as custom DNA oligo gBlock (Integrated DNA Technologies, IDT), containing the codon-optimized coding sequence with a FLAG-tag and cloned into pcDNA3.1. For PGAM5 lacking the C-terminal tail (ΔC), amino acids 1-277 of PGAM5 were subcloned into pcDNA3.1 inserting an early FLAG-tag followed by a stop codon. All constructs were verified by sequencing.

### Cell lines, transfection and RNA interference

Hek293T cells were grown in Gibco Dulbecco’s modified Eagle’s medium supplemented with 10% (v/v) fetal bovine serum at 37°C in 5% (v/v) CO2. For stable Hek293 T-REx cells, 1% (v/v) Gibco sodium pyruvate, 1% (v/v) Gibco GlutaMAX^TM^ (Thermo Fisher Scientific) and the required antibiotics 5 µg/mL blasticidine (Gibco) and 500 µg/mL geneticin-G418 (Gibco) were added additionally. Transient transfections were performed using 25 kDa linear polyethylenimine (Polysciences) (80) as had been described (28). If not otherwise indicated, 500 ng plasmid encoding PGAM5-FLAG and 300 ng plasmid encoding PARL were used.

Total transfected DNA (2 µg/well) was held constant by the addition of empty plasmid. If not otherwise stated, cells were harvested 36 h after transfection. For transfection of small interfering RNA (siRNA), 2x10^5^ Hek293 T-REx cells were seeded per well of a 6-well plate. After 24 h, cells were transfected with 20 nM siRNA-oligonucleotide, either Silencer™ Select NegCtrl#1 (4390843, Ambion) or OMA1 Silencer™ Select Pre-Designed siRNA (s41777, Ambion), using Lipofectamine RNAiMAX reagent (Thermo Fisher Scientific). After 48 h incubation with siRNA, cells were transfected with DNA as described above and harvested 24-36 h later. Knockdown was performed with 0.5 µg/mL doxycyclin for 6 days. Disruption of the mitochondrial membrane potential was achieved by incubating the cells with 10 µM CCCP from a stock in dimethylsolfoxid (DMSO) for 3 h. For inhibition of the proteasome, approx. 24 h pre-harvesting 2 μM MG132 (Calbiochem) were added from a 10,000x stock in DMSO. As a vehicle control, the same amount of DMSO was used for untreated samples. Cells were harvested and lysed in SDS-sample buffer.

### *in vitro* cleavage assay using purified proteins

MBP-PGAM5 expression, purification and the PARL cleavage assay were described before in (22).

### SDS-PAGE and western blotting

Proteins were resolved on Tris-glycine poly-acrylamide gels followed by western blot analysis. Transfected cells were solubilized in Tris-glycine SDS-PAGE sample buffer (50 mM Tris-Cl pH 6.8, 10 mM EDTA, 5% glycerol, 2% SDS, 0.01% bromphenol blue, 5% β- mercaptoethanol). All samples were incubated for 15 min at 65°C. Denaturated and fully-reduced proteins were resolved by Tris-glycine SDS-PAGE, blotted onto PVDF membrane (Immobilon-P, 0.45 μM pore size, Merck Millipore) via semi-dry blotting system and protein signal analyzed by using enhanced chemiluminescence to detect bound antibodies (WesternBright^TM^ ECL, Advasta). For detection, the ImageQuant™ LAS 4000 system (GE Healthcare) was used. Data shown are representative of at least three independent experiments. For quantification, we used the ImageJ software (http://rsb.info.nih.gov/ij/).

Statistical analysis was carried out using Prism 9.1.2 (226) software (GraphPad Software Inc.).

### Blue native PAGE of mitochondrial enriched crude membranes

If not indicated differently, all steps were performed on ice or at 4°C. Mitochondrial enriched crude membranes of Hek293T cells ectopically expressing PGAM5-FLAG were obtained by cell disruption followed by differential centrifugation. In brief, cells were detached by phosphate-buffered saline-EDTA and resuspended in isolation buffer (250 mM sucrose, 10 mM Tris-Cl pH 7.4, 10 mM HEPES pH 7.4, 0.1 mM EGTA, EDTA-free complete protease inhibitor cocktail (Roche Molecular Biochemicals). After 10 min incubation at 4°C, cells were lysed by passing six times through a 27-gauge needle. Cellular debris and nuclei were discarded after centrifugation at 200 xg for 5 min at 4°C. The supernatant was spun at 10,000 xg for 10 min at 4°C and the membrane pellet containing mitochondrial membranes was resuspended in isolation buffer and washed one more time. Further, the mitochondrial enriched crude membranes were solubilized with 1% Triton X-100 in (8 mM Tris-Cl pH 7.4, 20 mM NaCl, 0.6 mM MgCl2, 4% glycerol, 0.4 mM EGTA) supplemented with EDTA-free complete protease inhibitor cocktail (1xPI, Roche) and 1 mM PMSF. After removal of insoluble fraction by centrifugation at 14,000 rpm, supernatant was combined with a 1/40 volume of BN sample buffer (500 mM 6-aminohexanoic acid, 100 mM Bis-Tris pH 7.0, 5% Coomassie G250) before subjection onto Native-PAGE in self-casted Bis-Tris 6-20% acrylamide (AA-Bis, 40%, 32:1) gradient gels. Gels were run for 1 h at 150 V, buffer changed according to the manufacturers description and then continued at 230 V for 2-3 h.

Afterwards, gels were incubated for 15 min in BN-transfer buffer (25 mM Tris, 150 mM glycine, 0.02% SDS, 20% methanol) and were transferred at 100 mA for 70 min onto PVDF membrane (Immobilon-P, 0.45 μM pore size, Merck Millipore) using semi-dry blotting system. The PVDF membrane was incubated in fixation solution (40% methanol, 10% acetic acid), destained in methanol, washed in water, blocked in 5% milk TBS-Tween (10 mM Tris-Cl pH 7.4, 150 mM NaCl, 0.1% Tween 20), and analyzed using enhanced chemiluminescence (WesternBright^TM^ ECL, Advasta) by ImageQuant™ LAS 4000 system (GE Healthcare).

### Antibodies

The following antibodies were used at dilutions recommended by the manufacturer. For western blot analysis primary antibodies were used in 5% milk/TBS-T, secondary antibodies in TBS-T only: Mouse monoclonal anti-FLAG (M2) 1:1000 (F1804, Sigma-Aldrich), mouse monoclonal anti-PGAM5 1:1000 (CL0624, Thermo Fisher Scientific), rabbit polyclonal anti- PARL 1:300 (ab45231, Abcam), rabbit polyclonal anti-PARL 1:1000 (600-401-J27, Rockland), mouse monoclonal anti-β-Actin 1:3000 (A1978, Sigma-Aldrich), donkey anti-mouse igG (H+L) 1:10000 (715-035-150, Dianova), and donkey anti-rabbit igG (H+L) 1:10000 (711-035-152, Dianova). For immunofluorescence analysis (for method see below): Rabbit polyclonal anti-FLAG 1:500 (PA1-984B, Invitrogen) and mouse monoclonal anti- TOM20 1:400 (sc-17764, Santa Cruz Biotechnology), goat anti-mouse igG (H+L) Alexa Fluor 488 (A11029, Invitrogen) and goat anti-rabbit igG (H+L) Alexa 633 (A21070, Invitrogen).

### Immunofluorescent staining on fixed cells and microscopy

Hek293T cells were plated in 24-well plates on cover glass (Carl Roth) coated with poly-L-Lysine (Sigma-Aldrich). Cells were transfected with 125 ng of plasmid and total plasmid levels were adjusted to 500 ng with empty plasmid. For immunofluorescence analysis, cells were chemically fixed for 15 min with 4% formaldehyde (16% formaldehyde diluted in PBS, Thermo Scientific), washed 3x in PBS followed by permeabilization and blocking in PBS containing 0.1% Triton X-100 (EMD Millipore) and 20% fetal calf serum (TPBS-FCS) for 45 min. Subsequently, the fixed cells were probed with anti-TOM20 and anti-FLAG antibodies in TPBS-FCS for 1 h and washed 3x in PBS. After staining with fluorescently labelled secondary antibodies Alexa Fluor 488 and Alexa 633, both diluted in TPBS-FCS for 1 h, the slides were washed 3x in PBS, followed by Hoechst staining (1 µg/mL in PBS) for 10 min.

After washing 3x with PBS, the cover glasses were mounted with Fluoromount-G (Southern Biotech) on microscope slides. Samples were imaged with a LSM780 system (Carl Zeiss) using 405, 488, and 633 nm laser lines, a Plan-APOCHROMAT 63x 1.4NA oil objective (Carl Zeiss) and pinhole settings of 1AU with the Zeiss ZEN 2010 software. Image processing was performed using ImageJ (http://rsb.info.nih.gov/ij/).

### Liquid-state NMR

Unlabeled PGAM5 TM domain wt and mutant peptides were purchased from Core Unit Peptid-Technologien (University of Leipzig, Germany). For structure determination peptides were dissolved in 500 µL TFE-d2 and H2O (80:20) to a final concentration of 500 µM, pH was adjusted to 5.0. A set of homo- and heteronuclear liquid-state NMR spectra was acquired at 300 K on a 600 MHz Avance III spectrometer equipped with a CPTCI cryogenically cooled probehead (Bruker BioSpin, Germany). ^1^H, ^13^C and ^15^N resonances were assigned with ^1^H^1^H-TOCSY, ^1^H^13^C-HSQC and ^1^H^15^N-HSQC spectra at natural abundance. For structure calculation ^1^H^1^H-NOESY spectra were acquired with 200 ms mixing time. Data acquisition and processing was done with TopSpin (Bruker BioSpin, Germany) and CcpNMR Analysis was used for assignment and integration (81). Backbone dihedral angles were predicted based on chemical shift values with TALOS+ (82) and three-dimensional peptide structures were calculated with Aria2 (83) based on NOE derived distance restraints and dihedral angles. Graphical representations of the structures were created with PyMOL (The PyMOL Molecular Graphics System, ver. 2.3.4, Schrödinger, LLC).

### H/D exchange

For hydrogen-deuterium-exchange measurements dry peptides were dissolved in deuterated solvent, TFE-d3 and D2O (80:20), to a final concentration of 500 µM and two pD values, pD 4.0 and pD 5.0. A series of ^1^H^1^H-TOCSY spectra was acquired over a total period of 38 h.

Exchange rates were determined based on decreasing HNHα-cross-peak intensities with time.

### CD-spectroscopy

CD spectra were acquired on a JASCO J-810 spectrometer (Jasco, Pfungstadt, Germany) of IBG-2, KIT with 1 mm pathlength. Samples used for NMR measurements were diluted 10- fold to 50 µM peptide concentration. Scanning mode was set to 10 nm/min, scanning speed 8s, data pitch 1 nm and three spectra were accumulated. Measured was circular dichroism (CD), voltage (HT) and absorbance (Abs) from 180 to 250 nm. Data was analyzed using the BestSel online tool (84, 85).

### Data availability

All data is located in the manuscript. The atomic coordinates and experimental data used for structure calculation have been deposited in the Protein Data Bank (www.wwpdb.org) and BMRB (https://bmrb.io/). WT: 7QAM, 34681; C12L: 7QAL, 34680; C12S: 7QAO, 34682;

G17L: 7QAP, 34683. Structure statistics of PGAM5 wt and the three mutants can be found in the supplemental information.

## Acknowledgements

We thank the lab of M. Joanne Lemieux (University of Alberta, Canada) for the supply with yeast-purified recombinant human PARL, Alireza Pouya for help with setting up the microscopy analysis, and Thomas Langer for feedback to the manuscript. This work was supported by the grant Le2749/1-2 and MU1606/6-2 of the Deutsche Forschungsgemeinschaft (German Research Foundation) as part of 263531414/FOR2290. CMG acknowledges funding by the Helmholtz-Society. MS was supported by a Carl-Zeiss fellowship.

## Author contributions

Experiments were devised by VS, MS, MKL and CMG. VS conducted human cell-based gain- and loss-of-function assays, MBP-PGAM5 expression and *in vitro* cleavage assays. MS conducted CD-spectroscopy, liquid-state NMR, H/D exchange. EH conducted immunofluorescence microscopy analysis. The manuscript was written by VS and MKL, and edited by VS, MS and CMG.

## Supplemental Figures

**Figure S1.**
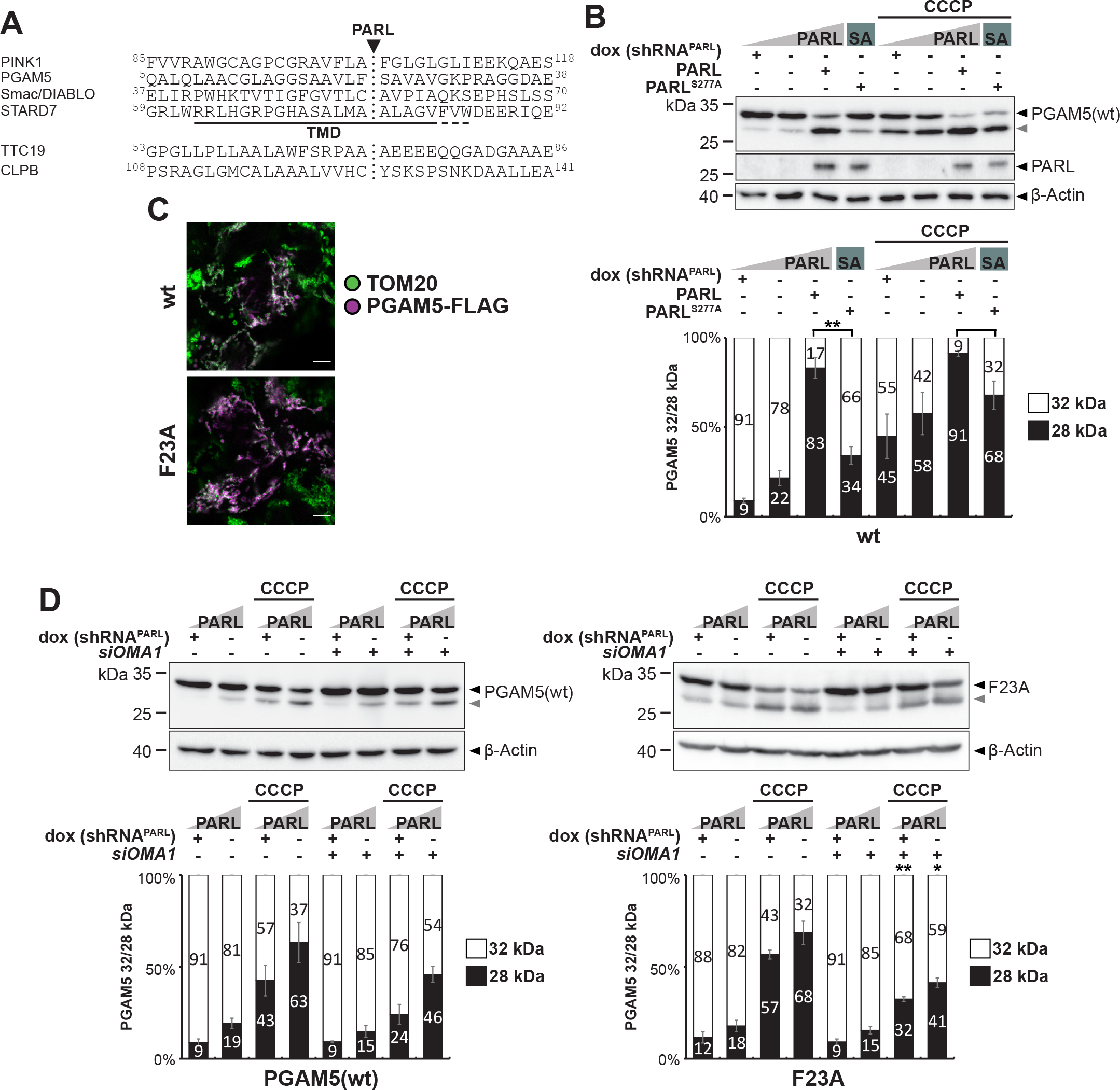
Bulky residue in P1 position shows only modest influence on PGAM5 processing. **(A)** TM domain (TMD) and cleavage site region (marked by dotted line) of so far identified PARL substrates (17,28,29,79,87,88). **(B)** PGAM5 processing analyzed in cell-based PARL gain- and loss-of-function assay as in Fig. 1B. Ectopic expression of PARL wt but not the catalytic-inactive PARL^S277A^, leads to increased processing. PGAM5 cleavage was stimulated by treating cells with the mitochondrial uncoupling agent CCCP. Grey arrowhead: 28 kDa cleavage fragment. Lower panel shows quantification of PGAM5 32/28 kDa distribution (n = 3, means ± SEM). Significant changes comparing PARL and PARL^S277A^ overexpression without and with CCCP treatment are indicated with black stars (*p ≤ 0.05, **p ≤ 0.01; unpaired two-tailed t-test). **(C)** Immunofluorescence analysis examines mitochondrial targeting of ectopically expressed PGAM5-FLAG constructs (purple) co-stained with endogenous TOM20 (green). Scale bar, 5 μm. **(D)** For *OMA1* knockdown, cells were transiently transfected with either a non-targeting siRNA or an *OMA1*-specific siRNA for 48 hours before transient transfection of the PGAM5-FLAG contructs. PGAM5 cleavage was stimulated by treating cells with CCCP. Grey arrowhead: 28 kDa cleavage fragment. Lower panels each show quantification of PGAM5 32/28 kDa distribution (n = 3, means ± SEM). Significant changes versus cells transfected with non-targeting siRNA are indicated with black stars (*p ≤ 0.05, **p ≤ 0.01; unpaired two-tailed t-test).

**Figure S2.**
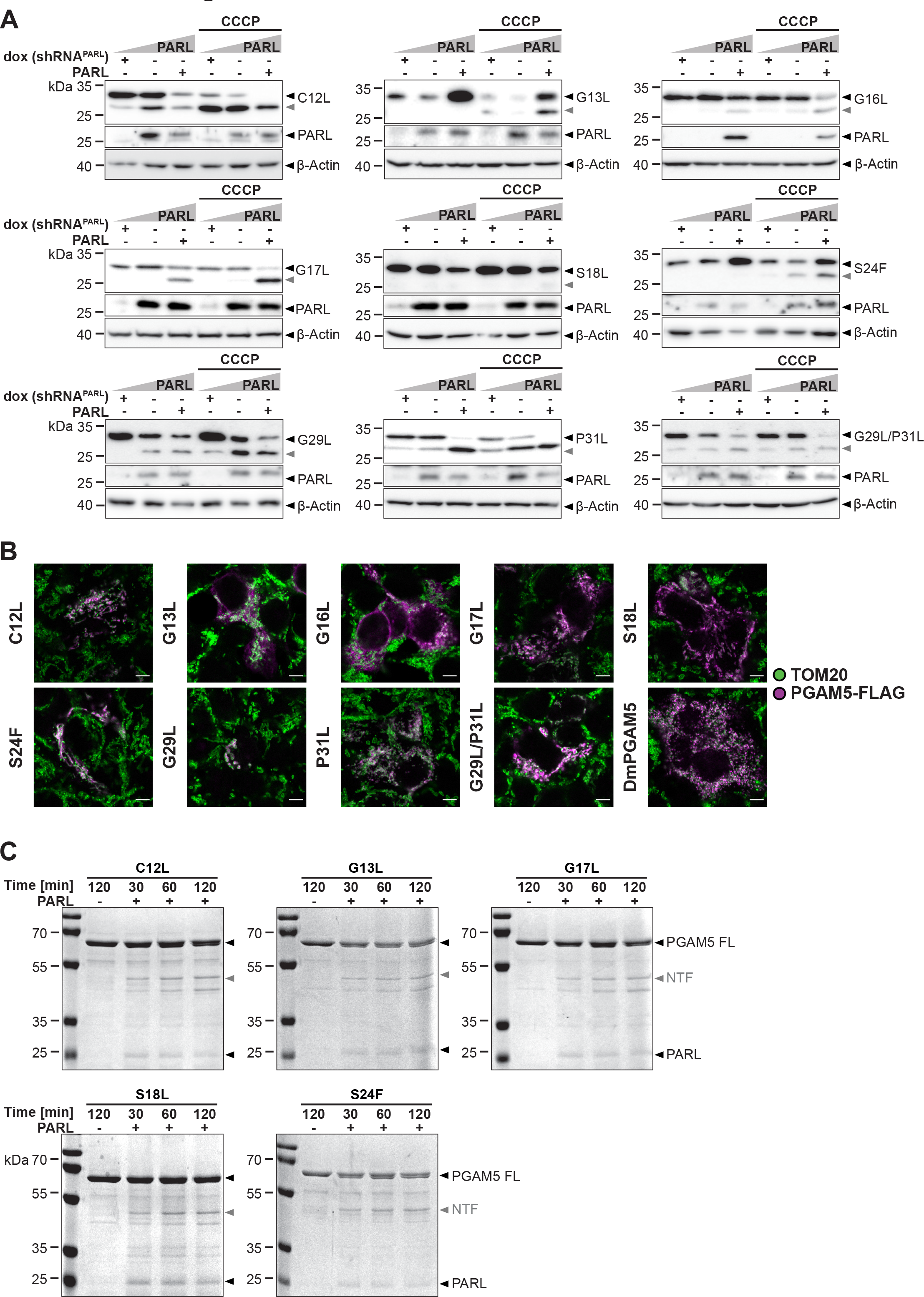
PARL-catalyzed cleavage of PGAM5 is influenced by multiple TM residues. **(A)** Western blot analysis of PGAM5 processing in a cell-based PARL gain- and loss-of-function assay as in Fig. 1B. Grey arrowhead: 28 kDa cleavage fragment. See Fig. 2B for quantification. **(B)** Immunofluorescence analysis examines mitochondrial targeting of ectopically expressed mutant PGAM5-FLAG constructs (purple) co-stained with endogenous TOM20 (green). Scale bar, 5 μm. **(C)** Incubation of detergent-solubilized and purified recombinant PARL with MBP-PGAM5 leads to generation of an N-terminal cleavage fragment (NTF) as resolved by SDS-PAGE and staining with Coomassie blue. PARL- dependent alternative cleavage fragments appeared as side-effects of the detergent background. FL: MBP-PGAM5 full length. See Fig. 2C for quantification.

**Figure S3.**
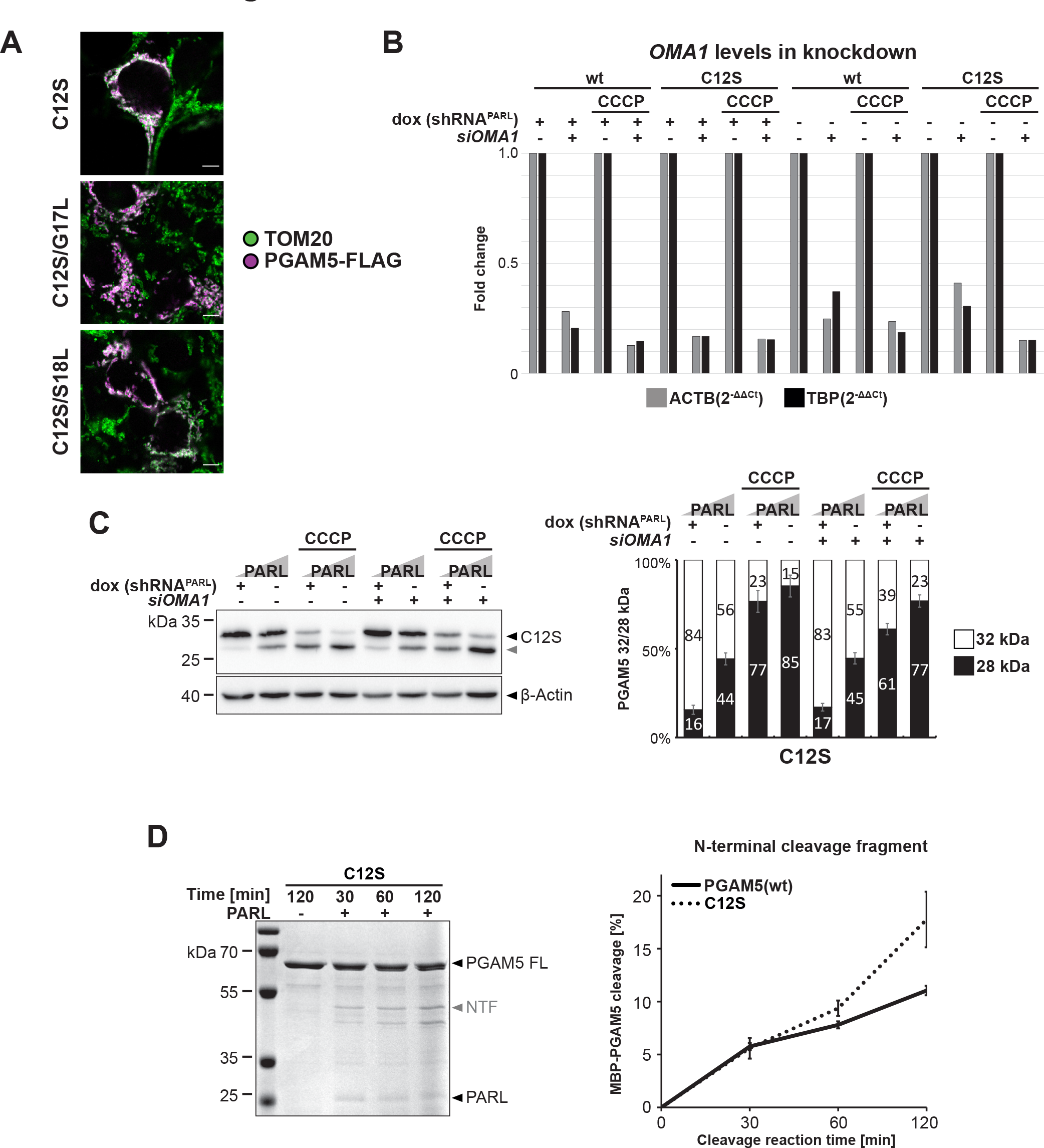
N-terminal substrate feature in PGAM5 important for PARL-catalyzed cleavage. **(A)** Immunofluorescence analysis examines mitochondrial targeting of ectopically expressed mutant PGAM5-FLAG constructs (purple) co-stained with endogenous TOM20 (green). Scale bar, 5 μm. **(B)** *OMA1* levels are knocked down by transient transfection of *OMA1*-specific siRNA. Normalization of relative gene expression by comparison to the reference genes β-Actin (ACTB) and TATA-box binding protein (TBP). **(C)** For *OMA1* knockdown, cells were transiently transfected with either a non-targeting siRNA or an *OMA1*-specific siRNA for 48 hours before transient transfection of the PGAM5-FLAG contructs. PGAM5 cleavage was stimulated by treating cells with CCCP. Grey arrowhead: 28 kDa cleavage fragment. Right panel shows quantification of PGAM5 32/28 kDa distribution (n = 3, means ± SEM). No significant changes versus cells transfected with non-targeting siRNA were observed (unpaired two-tailed t-test). **(D)** Incubation of recombinant PARL with MBP-PGAM5^C12S^ leads to more efficient generation of the N-terminal cleavage fragment when compared with the wt construct (n = 3, means ± SEM; see Fig. 1C for comparison). PARL-dependent alternative cleavage fragments appeared as side-effects of the detergent background. FL: MBP-PGAM5 full length.

**Figure S4.**
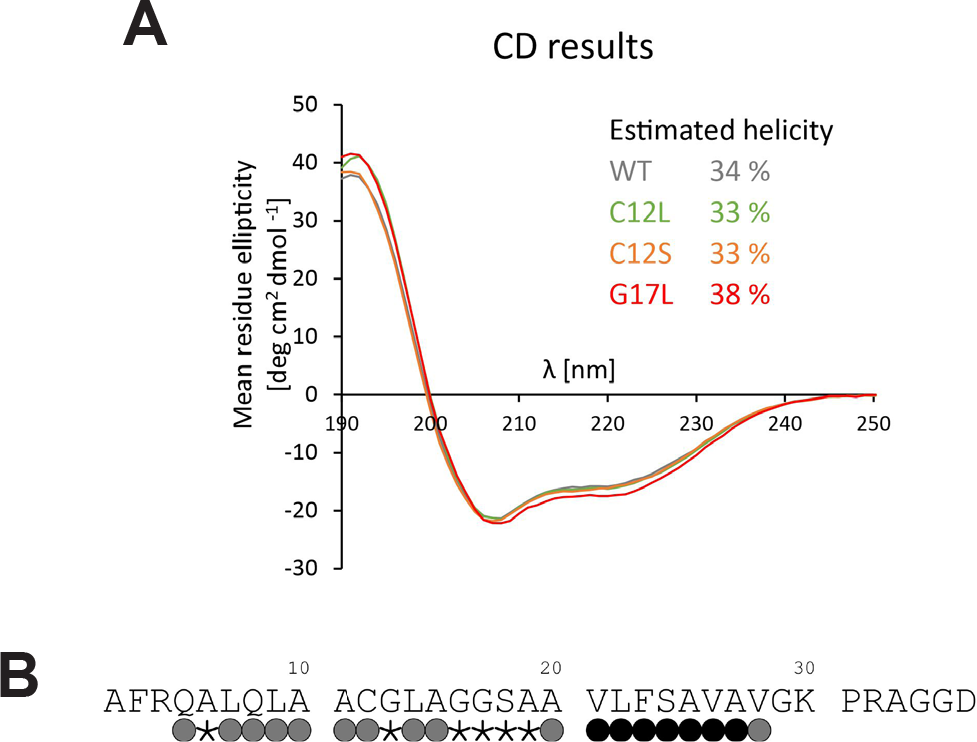
Structural properties of the PGAM5 TM domain. **(A)** CD results of wt and three mutants. Values are scaled to wt values. **(B)** Hydrogen-deuterium exchange of wt TM domain shows stable H-bonds directly before the cleavage site and at the N-terminus between Q8 an C12. Hydrogen bonds at the helix termini and between G13 and G17 are significantly weakened. Black dots indicate fast exchange, grey dots slow exchange. Some exchange rates could not be determined due to peak overlap, these residues are marked by an asterisk.

**Figure S5.**
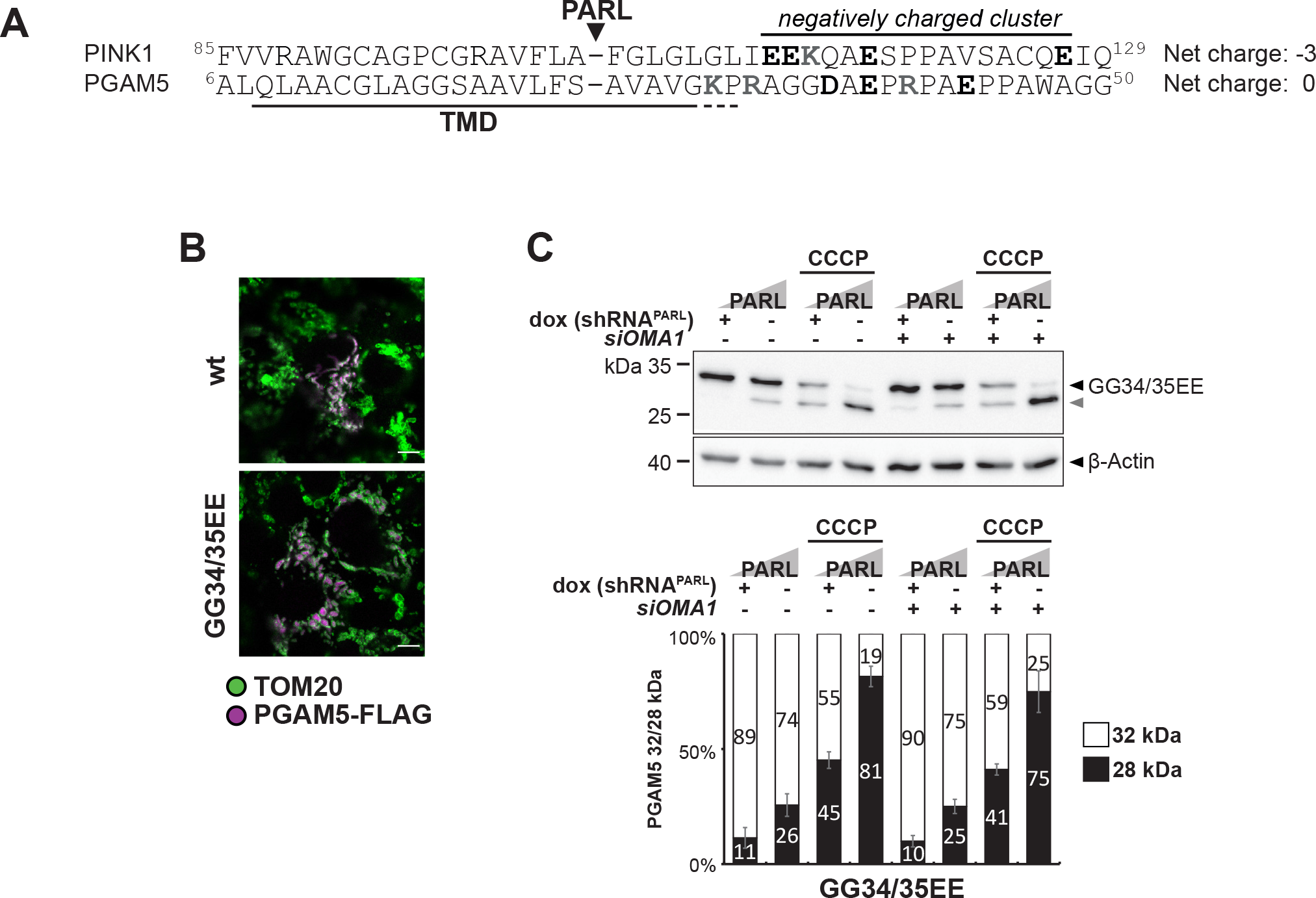
Negative charges in the PGAM5 juxtamembrane region influence cleavage efficiency under CCCP. **(A)** Negatively charged cluster in juxtamembrane region of PINK1 but not PGAM5. Net charge in juxtamembrane region of negatively charged amino acids (black) and positively charged amino acids (grey). **(B)** Immunofluorescence analysis examines mitochondrial targeting of ectopically expressed mutant PGAM5-FLAG constructs (purple) co-stained with endogenous TOM20 (green). Scale bar, 5 μm. **(C)** For *OMA1* knockdown, cells were transiently transfected with either a non-targeting siRNA or an *OMA1*-specific siRNA for 48 hours before transient transfection of the PGAM5-FLAG contructs. PGAM5 cleavage was stimulated by treating cells with CCCP. Grey arrowhead: 28 kDa cleavage fragment. Below the quantification of PGAM5 32/28 kDa distribution (n = 3, means ± SEM). No significant changes versus cells transfected with non-targeting siRNA were observed (unpaired two-tailed t-test).

**Figure S6.**
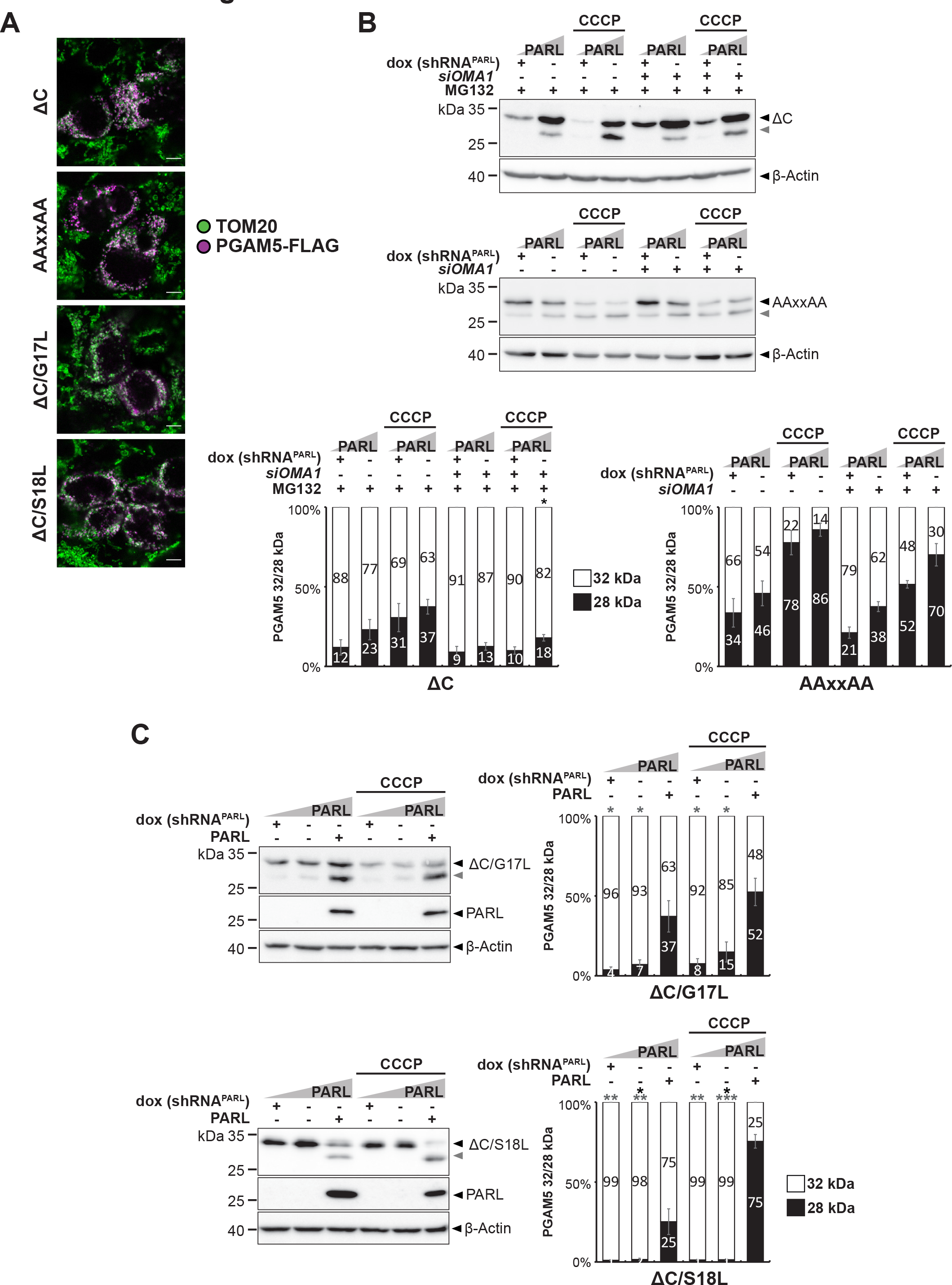
Formation of the PGAM5 higher order structure prevents PARL-catalyzed cleavage. **(A)** Immunofluorescence analysis examines mitochondrial targeting of ectopically expressed oligomerization-defecient mutant PGAM5-FLAG constructs (purple) co-stained with endogenous TOM20 (green). Scale bar, 5 μm. **(B)** For *OMA1* knockdown, cells were transiently transfected with either a non-targeting siRNA or an *OMA1*-specific siRNA for 48 hours before transient transfection of the PGAM5-FLAG contructs. PGAM5 cleavage was stimulated by treating cells with CCCP. Grey arrowhead: 28 kDa cleavage fragment. Lower panels show quantification of PGAM5 32/28 kDa distribution (n = 3, means ± SEM). No significant changes versus cells transfected with non-targeting siRNA were observed (unpaired two-tailed t-test). **(C)** Corresponding analysis of monomeric PGAM5 double mutants additionally containing TM domain mutations G17L and S18L (PGAM5^G17L^ΔC, PGAM5^S18L^ΔC). Grey arrowhead: 28 kDa cleavage fragment. Right panel shows quantification of PGAM5 32/28 kDa distribution (n = 3, means ± SEM). Significant changes versus wt PGAM5-FLAG are indicated with black stars, significant changes versus PGAM5^ΔC^-FLAG are indicated with grey stars (*p ≤ 0.05, **p ≤ 0.01, ***p ≤ 0.001; unpaired two-tailed t-test).

**Figure S7.**
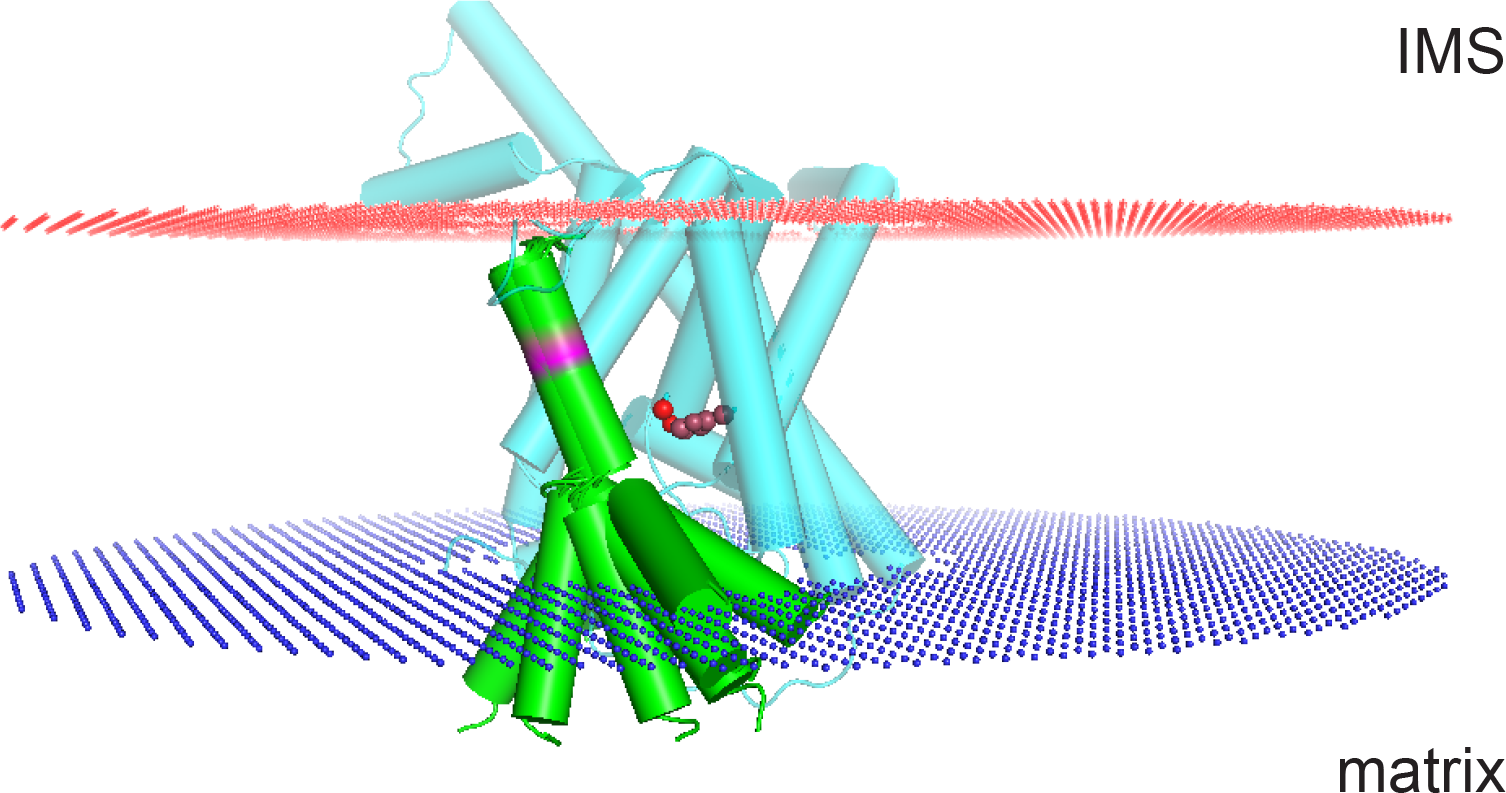
Hypothetical model of the PGAM5 TM domain bound by a putative PARL exosite. Model of PARL generated by AlphaFold, entry Q96HS1. Catalytic S277 and H335 are depicted in red facing the water-filled cavity of PARL, which opens to the matrix. Insertion depth for PARL AlphaFold model and the helical PGAM5 TM domain examinded by NMR into the inner mitochondrial membrane was determined with the OPM server. The amphipathic helix of PGAM5 is shown here with a submerged orientation that allows the charged and hydrophilic sidechains to be placed within the lipid headgroup area. Cleavage site within the C-terminal helix of PGAM5 TM domain (F23-S24) is depicted in magenta. Red area: upper lipid layer towards mitochondrial IMS, blue area: lower lipid layer towards mitochondrial matrix.

**Table S1.** Structure statistics of PGAM5 WT TMD and four single point mutants. All values refer to the ensemble of 20 structures with the lowest energy from 400 calculated structures. NOE: Nuclear Overhauser Effect, RMSD: root-mean-square deviation of atomic

|  | WT | C12L | C12S | G17L |
| --- | --- | --- | --- | --- |
| **Total restraints used** | 269 | 279 | 261 | 274 |
| unambiguous NOE restraints | 269 | 279 | 261 | 274 |
| Intraresidue | 106 | 107 | 124 | 127 |
| Sequential ($\vert i-j\vert =1$) | 85 | 87 | 87 | 93 |
| Medium range ($1 < \vert i-j\vert < 4$) | 61 | 66 | 47 | 40 |
| Long range ($\vert i-j\vert \geq 4$) | 17 | 19 | 3 | 17 |
| Ambiguous NOE restraints | 0 | 0 | 0 | 0 |
| **Statistics for structure calculations** |  |  |  |  |
| RMSD of bonds (Å) | 0.001 +/- 0.00007 | 0.001 +/- 0.00007 | 0.001 +/- 0.00006 | 0.001 +/- 0.0001 |
| RMSD of bond angles (°) | 0.253 +/- 0.005 | 0.260 +/- 0.005 | 0.252 +/- 0.004 | 0.285 +/- 0.011 |
| RMSD of improper torsions (°) | 0.108 +/- 0.007 | 0.098 +/- 0.007 | 0.102 +/- 0.007 | 0.146 +/- 0.02 |
| **Final Energies (kcal mol<sup>-1</sup>)** |  |  |  |  |
| E<sub>total</sub> | -1044 +/- 30 | 1057 +/- 24 | -1034 +/- 30 | -1058 +/- 22 |
| E<sub>bonds</sub> | 0.369 +/- 0.064 | 0.360 +/- 0.059 | 0.403 +/- 0.056 | 0.566 +/- 0.121 |
| E<sub>angles</sub> | 8.35 +/- 0.33 | 8.98 +/- 0.32 | 8.24 +/- 0.26 | 10.90 +/- 0.89 |
| E<sub>impropers</sub> | 0.432 +/- 0.057 | 0.362 +/- 0.055 | 0.387 +/- 0.053 | 0.823 +/- 0.287 |
| E<sub>dihed</sub> | 134.0 +/- 0.98 | 134.2 +/- 0.81 | 133.3 +/- 0.82 | 140.2 +/- 1.78 |
| E<sub>vdW</sub> | -207.6 +/- 4.3 | -203.22 +/- 4.28 | -195.96 +/- 3.2 | -210.1 +/- 2.7 |
| E<sub>NOE</sub> | -980.1 +/- 28.7 | -988.25 +/- 25.5 | -981.0 +/- 29.5 | -1001 +/- 22 |
| **Coordinate precision (Å)** |  |  |  |  |
| RMSD of backbone (N,CA,C,O) of all residues | 3.06 | 4.47 | 4.20 | 4.35 |
| RMSD of all heavy atoms of all residues | 3.56 | 5.11 | 5.05 | 5.05 |
| RMSD of backbone (N,CA,C,O) of ordered residues (3:28) | 2.18 | 2.30 | 2.95 | 3.42 |
| RMSD of all heavy atoms of ordered residues (3:28) | 2.67 | 2.96 | 4.0 | 2.37 |

positions.

